# Data-driven spectral analysis for coordinative structures in periodic systems with unknown and redundant dynamics

**DOI:** 10.1101/511642

**Authors:** Keisuke Fujii, Naoya Takeishi, Benio Kibushi, Motoki Kouzaki, Yoshinobu Kawahara

## Abstract

Living organisms dynamically and flexibly operate a great number of components. As one of such redundant control mechanisms, low-dimensional coordinative structures among multiple components have been investigated. However, structures extracted from the conventional statistical dimensionality reduction methods do not reflect dynamical properties in principle. Here we regard coordinative structures in biological periodic systems with unknown and redundant dynamics as a nonlinear limit-cycle oscillation, and apply a data-driven operator-theoretic spectral analysis, which obtains dynamical properties of coordinative structures such as frequency and phase from the estimated eigenvalues and eigenfunctions of a composition operator. First, from intersegmental angles during human walking, we extracted the speed-independent harmonics of gait frequency. Second, we discovered the speed-dependent time-evolving behaviors of the phase on the conventional low-dimensional structures by estimating the eigenfunctions. Our approach contributes to the understanding of biological periodic phenomena with unknown and redundant dynamics from the perspective of nonlinear dynamical systems.

## 1. Introduction

Living organisms dynamically and flexibly operate a great number of components such as neurons, muscle fibers, and skeletal joints. These phenomena can be seemingly regarded as dynamical systems based on some specific rules; however, it is often intractable to find optimal solutions due to a number of combinations of controlled elements and environments. In the field of neuroscience or motor control, for example, this problem has been referred to as Bernstein’s degrees of freedom problem or redundant problem [1]. For this problem, studies in a forward modeling approach have accomplished forward simulations by introducing various bio-inspired systems (e.g., [2, 3, 4, 5, 6]), and by solving parameter optimization problems (e.g., [7, 8]). However, the dominance laws of a real organism behavior have sometimes been complicated or unclear. Recently, a backward or data-driven approach, which is expected to extract essential information from observed data, has attracted attention as an effective way to understand them.

One of the popular research subjects of the unknown and redundant dynamic phenomena is human locomotion, which can be performed at various speeds by solving the redundancy problem of many skeletal joints. This is because it seems to be based on specific rules (i.e., can be regarded as a cyclic movement) but the governing equation is not fully understood; thus, from a long time ago, it has attracted attention in many scientific and engineering fields such as neuroscience (e.g., [9, 10]), physics (e.g., [11, 12, 13]), clinical medicine (e.g., [14, 15, 16]), behavioral science (e.g., [17, 18]), robotics (e.g., [8, 5]), and pattern recognition (e.g., [19, 20]). For the redundant problem, some researchers have discovered synergistic, global, and coordinative structures among controlled components in joint motion (e.g., [10]) and muscle activities (e.g., [9]), which may contribute to solving the problem. In the early days, for locomotion, researchers have found that the three-dimensional trajectory of elevation angles of limb segments lies close to a two-dimensional plane [10, 21, 22], and have suggested that this planar law (or low-dimensional structure) may reflect intersegmental coordination. The planar law of intersegmental coordination has been observed at different walking speeds [23], forward or backward direction [24], and different levels of body weight unloading [25]. It has also been observed in cats [26], monkeys, and human infants [27]. The idea of the low-dimensional structures was also supported by the successful clinical application (e.g., [16]) and the accomplishment of walking model simulation [28]. However, since the conventional methods to extract coordinative structures have used statistical dimensionality reduction methods with assumptions of independence of sampling such as principal component analysis (PCA), the extracted structure does not reflect dynamical properties in principle and still, there has been little theoretical progress in this field. Therefore, an extraction method of the coordinative structures based on dynamical properties from data is needed. In other words, our motivation in this study is to understand the global dynamics behind the data obtained from human locomotion, of which governing equations are not fully known (e.g., neural inputs: for details, see Materials and Methods).

As a method to obtain a global modal description of nonlinear dynamical systems, operator-theoretic approaches have attracted attention such as in applied mathematics, physics and machine learning. One of the approaches is based on the composition operator (usually referred to as the Koopman operator [29, 30]), which defines the time evolution of observation functions in a function space, rather than directly defines the time evolution in a state space from a classical and popular view of the analysis (e.g., [31]). The advantage to use the operator-theoretic approach is to lift the analysis of nonlinear dynamical systems to a linear (but infinite-dimensional) regime, which is more amenable to the subsequent analysis. For example, spectral analysis of the Koopman operator can obtain dynamical properties such as frequency with delay/growth rate and coordinative (or coherent) structures corresponding to these properties. Among several estimation methods, one of the most popular algorithms for spectral analysis of the Koopman operator is dynamic mode decomposition (DMD) [32, 33]. The benefit of DMD is to extract such a global modal description of a nonlinear dynamical system from data, unlike other unsupervised dimensionality reduction methods such as PCA or singular value decomposition (SVD) for static data. DMD has been successfully applied in many real-world problems, such as analyses of epidemiology and electrocorticography (e.g., [34, 35]). The extracted coordinative structure based on the above dynamical properties by DMD is called *DMD modes*. Among many variants of DMD (for details, see Materials and Methods), Hankel DMD [36] theoretically yields the eigenvalues and eigenfunctions of a Koopman operator (called *Koopman eigenvalues* and *eigenfunctions*) by applying DMD to a Hankel matrix (or delay embedding matrix). The eigenfunction, which provides linearly evolving coordinates in the underlying state space [37], can describe the phase in (asymptotically) stable dynamical systems such as a classical Van del Pol oscillator model [38, 36]. Therefore, the first contribution of our study is to extract dynamical information such as a frequency, phase, and its coefficients (i.e., coordinative structures) for the understanding of biological periodic systems with redundant and (partly) unknown governing equations.

Furthermore, from the perspective of phase reduction, which has been studied in mathematics (e.g., [39]) and biology (e.g., [40]), we can also regard the underlying coordinative structure as a dynamical system with a limit cycle and reduce a periodic system to a phase model using our approach. In phase reduction, a limit-cycle oscillator (possibly evolving in high-dimensional space) is approximated to a phase oscillator (evolving on a one-dimensional circle) which is more amenable to mathematical analysis. Phase reduction has been used to analyze such as neural network models [41, 42, 43] and a circadian gene network model [44], but there have been fewer direct applications to the actual biological systems with unknown governing equations. Other researchers [45] reviewed the legged locomotion dynamics including a phase reduction model of neural oscillators, but that of an actual biological (e.g., intersegmental) motion was unknown. Therefore, the second contribution of our study is to obtain a phase reduction model in periodic movements of living organisms with unknown governing equations by a data-driven, operator-theoretic spectral analysis. Compared with the conventional approaches for collective motion dynamics with unknown governing equations (e.g., using a DMD variant [46]), we adopt Hankel DMD, which has strong theoretical support to understand the biological systems as dynamical systems or a phase model. In this study, we confirmed that the theoretical requirements are satisfied for locomotion data from the viewpoint of the convergence of estimation error. Although other locomotion studies obtained phase descriptions using local joint angles [47, 48, 49, 13] or a global one [50], in this study, we obtain global phase descriptions with various frequencies from locomotion data at various speeds. Furthermore, we examine the difference and similarity between the conventional and our approaches by investigating the extracted dynamical information on the conventional coordinative structures.

The purpose of this research is to clarify the dynamical properties of coodinative structures in periodic data with (partly) unknown and redundant dynamics by a data-driven spectral analysis of nonlinear dynamical system called Hankel DMD. The remainder of this paper is organized as follows. In Results section, we present the results of an application of (Hankel) DMD on actual human locomotion and for validation, on the simulation data of a double pendulum (has explicit analytical solutions) and a walking model (has explicit governing equations). In Discussion section, we follow up with discussions and conclusions. Finally, we describe Materials and Methods including the experimental setups, the background of Koopman spectral analysis, various DMDs including Hankel DMD, and the simulation setups.

## 2. Results

### Decomposition of segmental angles

First, we show the definition and an example of segmental angles human locomotion in Fig 1a and b, and show representative examples of intersegmental coordination by five decomposition methods in Fig 2: the conventional decomposition method using SVD (we call it SVD-based method), basic two DMDs, and two Hankel DMDs for each row. In the analysis, we used three elevation angles of the right thigh, shank and foot in a one cycle during 2.0 km/h walk for a participant (*T* = 126 [frame]), according to the previous studies such as [10] (for other experimental setups, see Materials and Methods). SVD-based method in the first row of Fig 2 shows similar two principal components in the previous study [51], in which the first component in Fig 2c seems to relate to limb orientation (approximately one cycle per one gait cycle) and the second component in Fig 2d seems to relate to limb length (roughly two cycles per one gait cycle). For evaluation, we used variability accounted for (VAF) and reconstruction error which are commonly used in non-orthogonal dimensionality reduction methods (for details, see Materials and Methods). VAF was also similar to the previous work in which on average the first two principal components accounted for more than 99% of the data (the mean reconstruction error for all walking speeds in three components is 0.0243 ± 0.0043).

**Figure 1:**
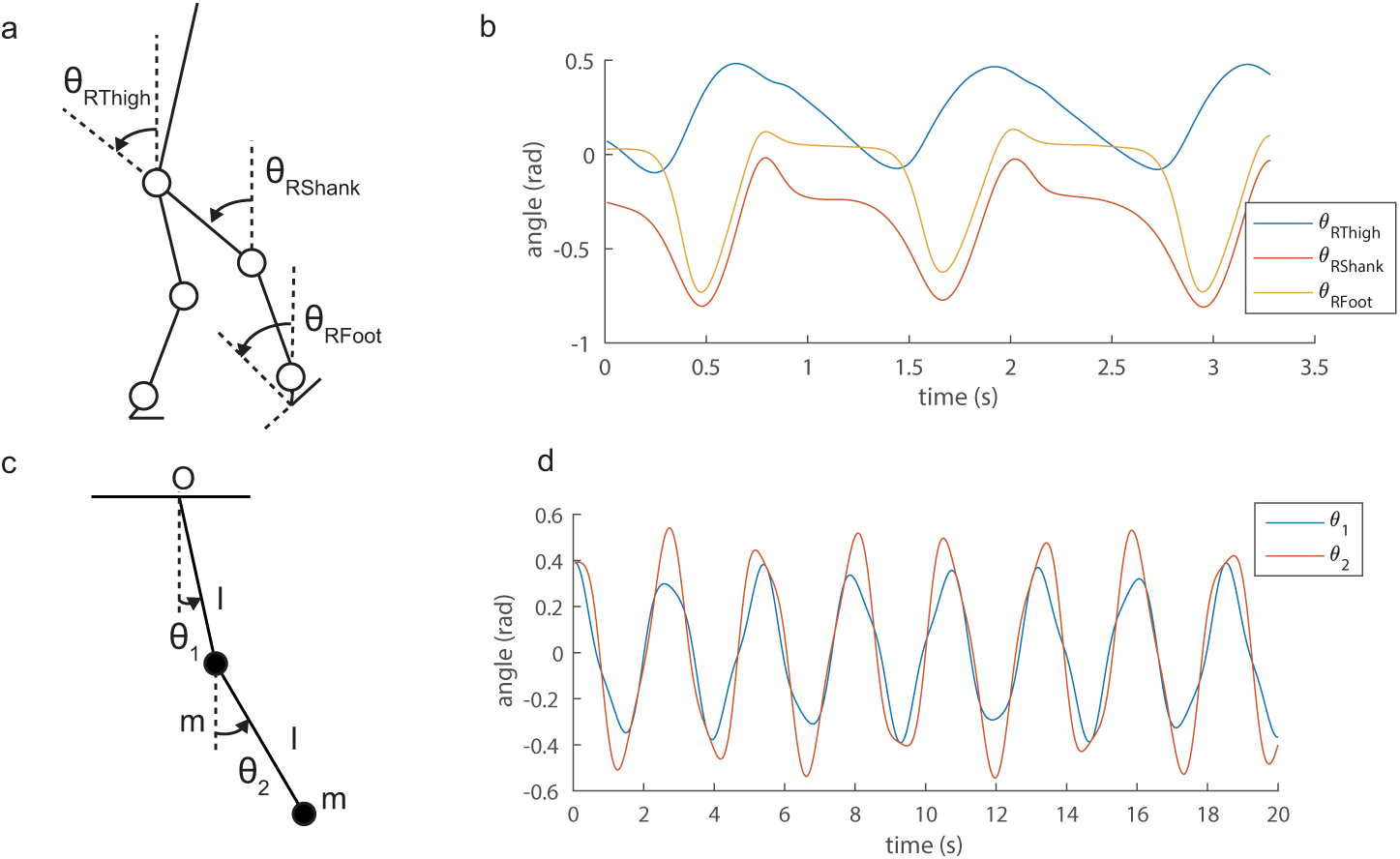
Angle definitions and examples of angle time-series. Definition of angles of human body (a) and double pendulum (c) are shown. Examples of angle time-series for (b) human locomotion and (d) double pendulum are indicated.

**Figure 2:**
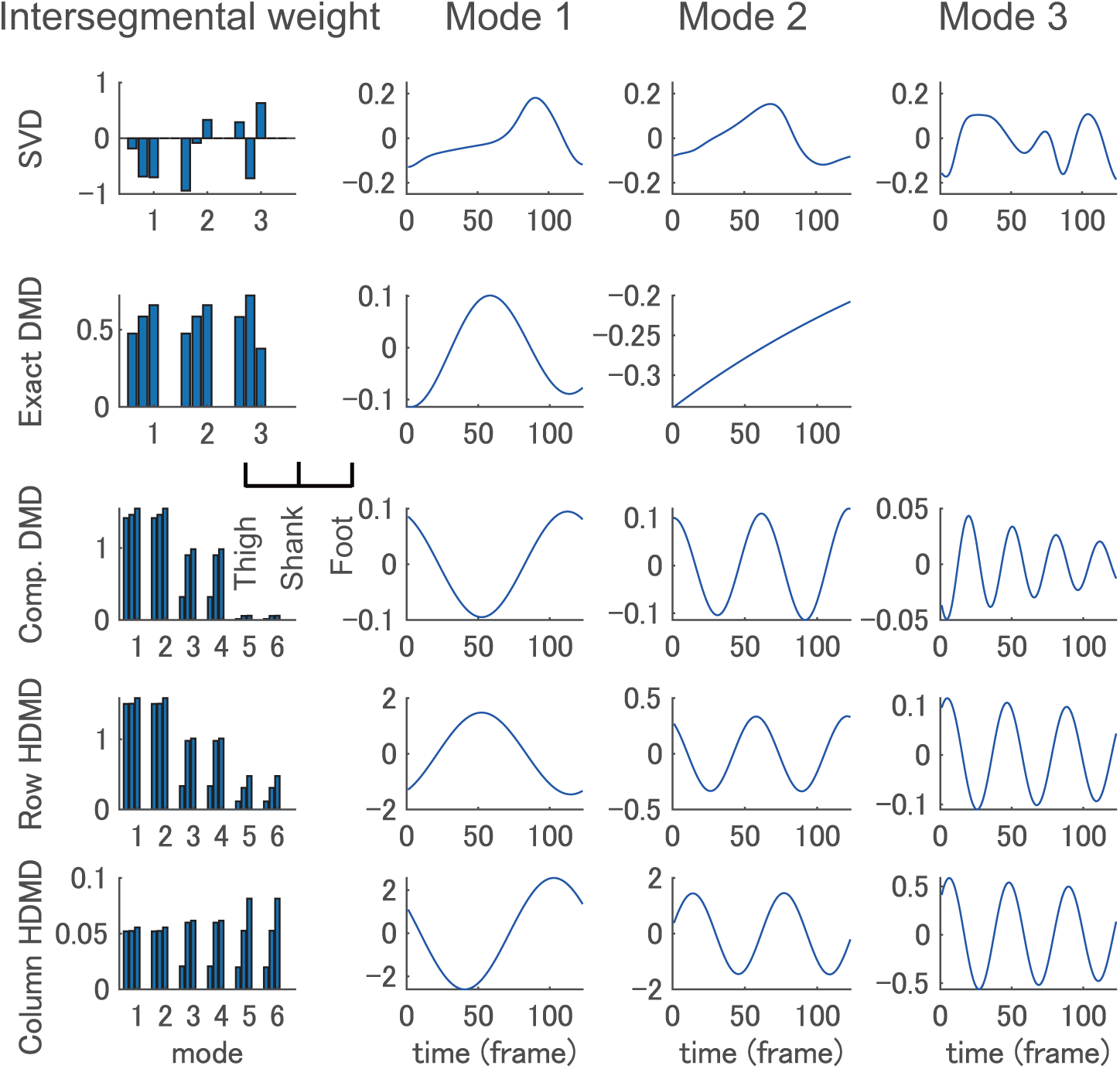
Representative examples of five decomposition methods. Examples of decomposition results into intersegmental weights (or DMD modes) in column 1 and time dynamics in the other columns by conventional SVD-based method (row 1), exact DMD (row 2), companion-matrix DMD (row 3), row-type Hankel DMD (row 4), and column-type Hankel DMD (row 5) during a 2.0 km/h walk are shown. SVD-based method shows similar two principal components in the previous study [51]. Exact DMD shows that only two conjugate and one modes because of the data dimension *d* = 3. In companion-matrix DMD and the two Hankel DMDs, we show only three pairs of conjugate modes that show the highest VAF. Note that all DMDs compute complex-valued DMD modes and time dynamics, but here we only illustrate the real part of them. VAF and reconstruction error of the dominant modes were higher and lower for SVD-based method, column-type Hankel DMD, row-type Hankel DMD, companion-matrix DMD, and exact DMD in this order.

Next, we explain the examples of the results of various DMDs. For the background and drawbacks of two basic DMDs (exact DMD and companion-matrix DMD), and the procedures of Hankel DMDs, see Materials and Methods. We compute row- and column-type Hankel DMDs for two different computational procedures. The example of exact DMD in row 2 of Fig 2 using the same data matrix in SVD-based method shows that only two conjugate and one modes because of the constraint of data dimension *d* = 3. Note that all DMDs compute complex-valued DMD modes and time dynamics, but here we only show the real part of them. Obviously from a theoretical perspective, companion-matrix DMD and the two Hankel DMDs from rows 3 to 5 of Fig 2 had more modes than exact DMD. Here, for clarity, we show only three pairs of conjugate modes which show the highest VAF in this order. Although all four DMDs extracted a similar pair of temporal dynamics with approximately one cycle (in column 2 of Fig 2), companion-matrix DMD and the two Hankel DMDs also extracted those with approximately two and three (or four) cycles. Regarding VAF and reconstruction error of the dominant modes, column-type Hankel DMD shows the best performance among the DMDs. On average among all walking speeds and participants, VAF was – 9.76 ± 6.35%, 26.70 ± 16.94%, 76.13 ± 0.59%, and 76.18 ± 0.48%, and reconstruction error was 0.3485 ± 0.0597, 0.2891 ± 0.0716, 0.0393 ± 0.068, and 0.0365 ± 0.074 for exact (a pair of the modes), companion-matrix, row-type and column-type Hankel DMDs (three pairs), respectively. We further evaluate the differences in the next subsection.

For DMD modes, in column 1 of Fig 2, those were similar among three DMDs in larger weight of foot angle than those of thigh and shank angles. Note that all DMDs compute complex-valued DMD modes and time dynamics, but here we only illustrate the real part of them. As shown in Fig 3, this tendency was consistent among various walking speeds in row-type Hankel DMD (the results of column-type Hankel DMD are shown in Fig S8). As a dynamical system to obey explicit governing equations, the results of the double pendulum and walking model simulation data are shown in Figs S3 and S4, respectively. Note that although in reconstruction error SVD obviously outperformed any DMD method, DMD methods can extract dynamical information that SVD cannot obtain, as shown in the below subsection.

**Figure 3:**
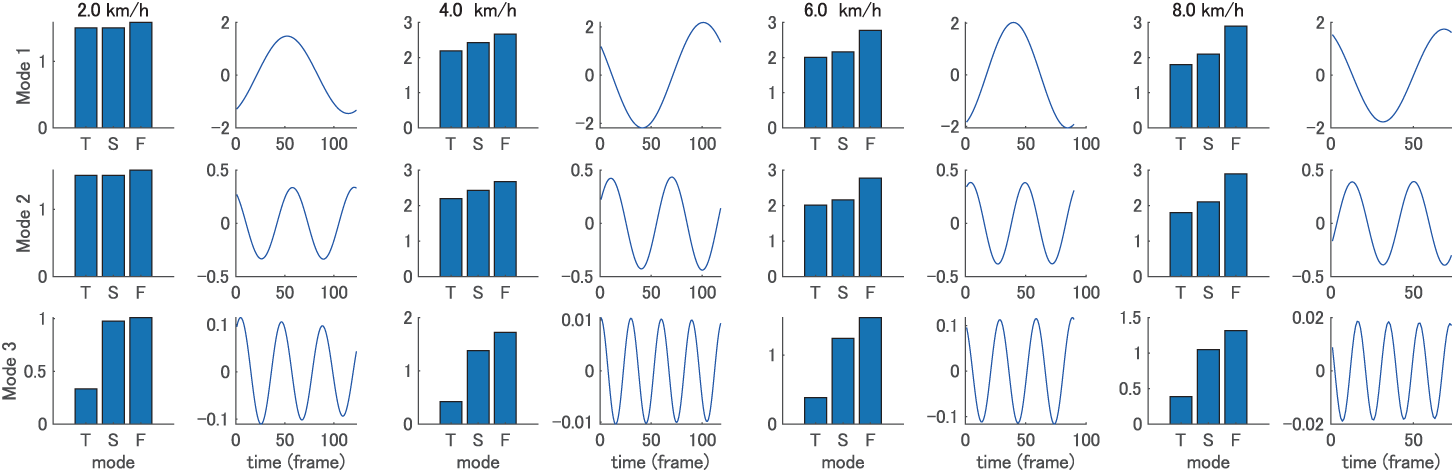
Representative examples of row-type Hankel DMD modes at various sppeds. Examples of three dominant row-type Hankel DMD modes (each left bar) and time dynamics (each right series) during a 2.0 to 8.0 km/h walk (from left to right in this order) are shown. T, S, and F indicate thigh, shank, and foot angles, respectively. Note that we show three modes with similar time dynamics (but different gait cycles and frequencies). The DMD modes were similar among four velocities in terms of the larger weight of foot angle than that of thigh and shank angles.

### Convergence and embedding dimension of Hankel DMDs

Next, we quantitatively validated that Hankel DMDs are applicable to human walking data. To compute both Hankel DMDs, we need to determine the dimension *p* of truncated SVD (or the number of Koopman eigenvalues) and the dimension *m* of delay embedding (for details, see Materials and Methods). Theoretically, Hankel DMDs with sufficient *m* obtain Koopman eigenvalues and eigenfunctions for a limit cycle [36]. As will be mentioned in Materials and Methods, we selected the dimensions *m* and *p* by considering the convergence and the avoidance of fitting to higher-frequency dynamics. First, from the perspective of the convergence, we investigated the sufficient *m* and *p* in Fig 4 for three representational walking speeds of a participant. We used the convergence of the average reconstruction errors among participants to investigate the convergence of the estimation error of Koopman eigenvalues and modes. Overall, both column- and row-type Hankel DMDs with larger *m* and *p* converged to an error < 0.01. In particular, the effect of *m* seemed to depend on walking speed (a higher speed or shorter gait cycle converged faster with an increase of *m*), but that of *p* did not seem to be independent of the speed if *m* is sufficient. For example, as a guide, if we select *p* = 50 and *m* = *T* or 2*T* for column- and row-type Hankel DMDs (for *T*, see Fig 4a-c top), the average error for all velocities and participants converged to 0.0013 ± 0.0006 and 0.0056 ± 0.0098 in column- and row-type Hankel DMDs, respectively.

**Figure 4:**
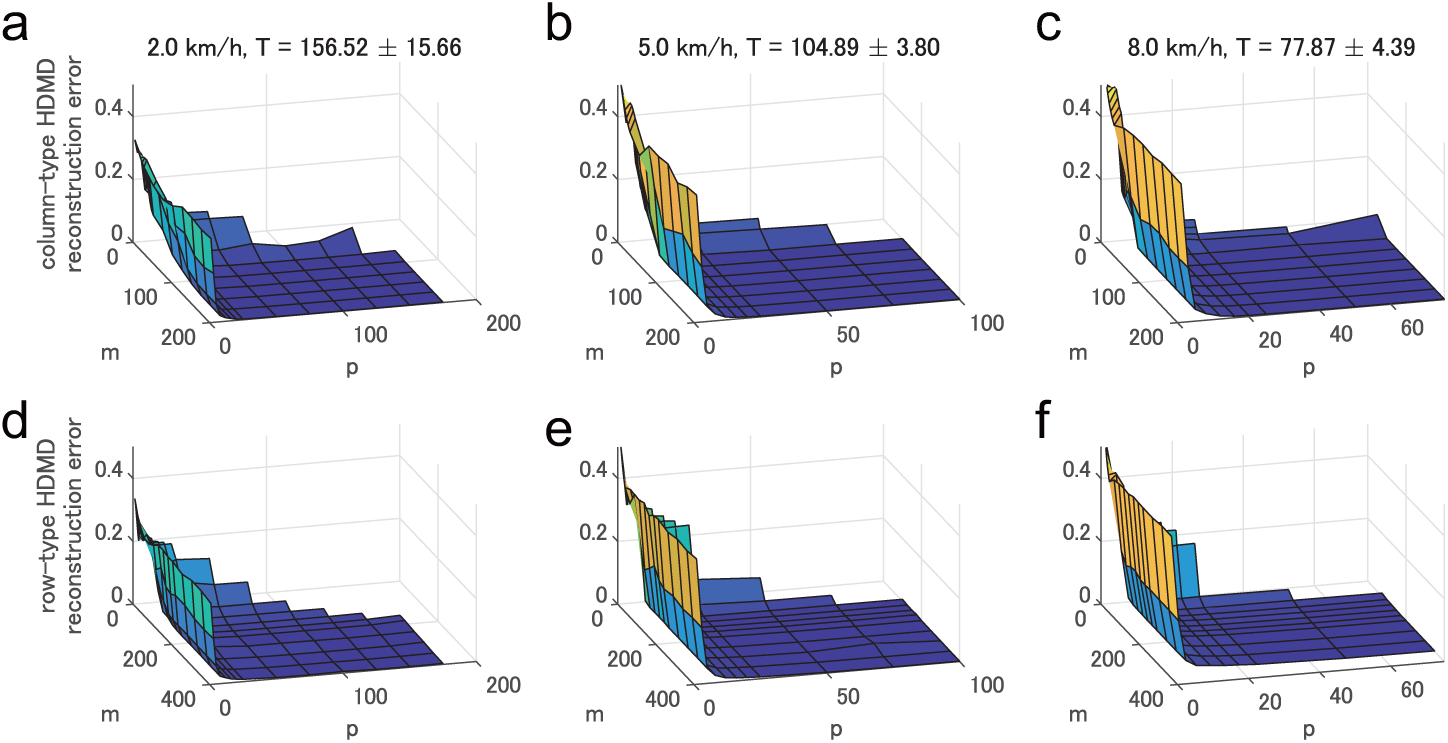
Convergence of Hankel DMDs. Examples of the reconstruction error for various *m* and *p* for column-type (a-c) and row-type (d-f) Hankel DMDs during 2.0 km/h (a and d), 5.0 km/h (b and e), and 8.0 km/h (c and f) walk are shown. Overall, both column- and row-type Hankel DMDs with larger *m* and *p* converged to a certain error. In particular, the effect of *m* seemed to depend on walking speed (i.e., a higher speed or shorter gait cycle converged faster with an increase of *m*), but that of *p* did not seem to be independent of the walking speed if *m* is sufficient. We selected *p* = 50 and *m* = *T*, 2*T* for column- and row-type Hankel DMDs. One gait cycle for each velocity is shown in (a-c) top.

Second, from the viewpoint of the avoidance of fitting to high-frequency dynamics, it may be better to select *m* and *p* as small as possible while satisfying the condition that the error converges. Therefore, we selected *m* and *p* as the above values (*p* = 50 and *m* = *T* or 2*T*). As a dynamical system to obey explicit governing equations, the results of similar convergence for the double pendulum (known frequency) in Fig 1c and d, and walking model simulation data (unknown frequency similar to human data) are shown in Fig S2.

### DMD (Koopman) eigenvalues

Next, we show that Hankel DMDs can extract the dynamical properties regarding temporal frequencies behind intersegment angles during locomotion. First, we explain it using representative distributions of eigenvalues and temporal frequency spectra computed by exact DMD, companion-matrix DMD, and column- and row-type Hankel DMDs in Fig 5 (we used the same example of Fig 2). For all DMDs, all eigenvalues were almost on the unit circle; however, the distributions were different. Both types of Hankel DMD in Fig 5e and g with the dimension of the truncation *p* = 50 show a similar distribution (again, *p* is the number of eigenvalues) whereas the eigenvalues of companion-matrix DMD in Fig 5c (*p* = *T* = 126 without truncation) uniformly occupied on the unit circle and those of exact DMD in Fig 5a (*p* = *d* = 3 without truncation) were near the real axis. Next, in temporal frequency domain, we show DMD and Fourier spectra averaging among all data dimensions of the input matrix. Although exact DMD in Fig 5b extracted low-frequency component among Fourier spectra, other DMDs obtained five or more harmonic frequencies (shown as light blue dashed line) of the gait frequency. In particular, row-type Hankel DMD in Fig 5h can extract similar spectra to the averaged Fourier spectra, whereas the higher-frequency norms of companion-matrix DMD in Fig 5d decayed in higher frequencies and the spectra of column-type Hankel DMD in Fig 5f were relatively different from the averaged Fourier spectra.

**Figure 5:**
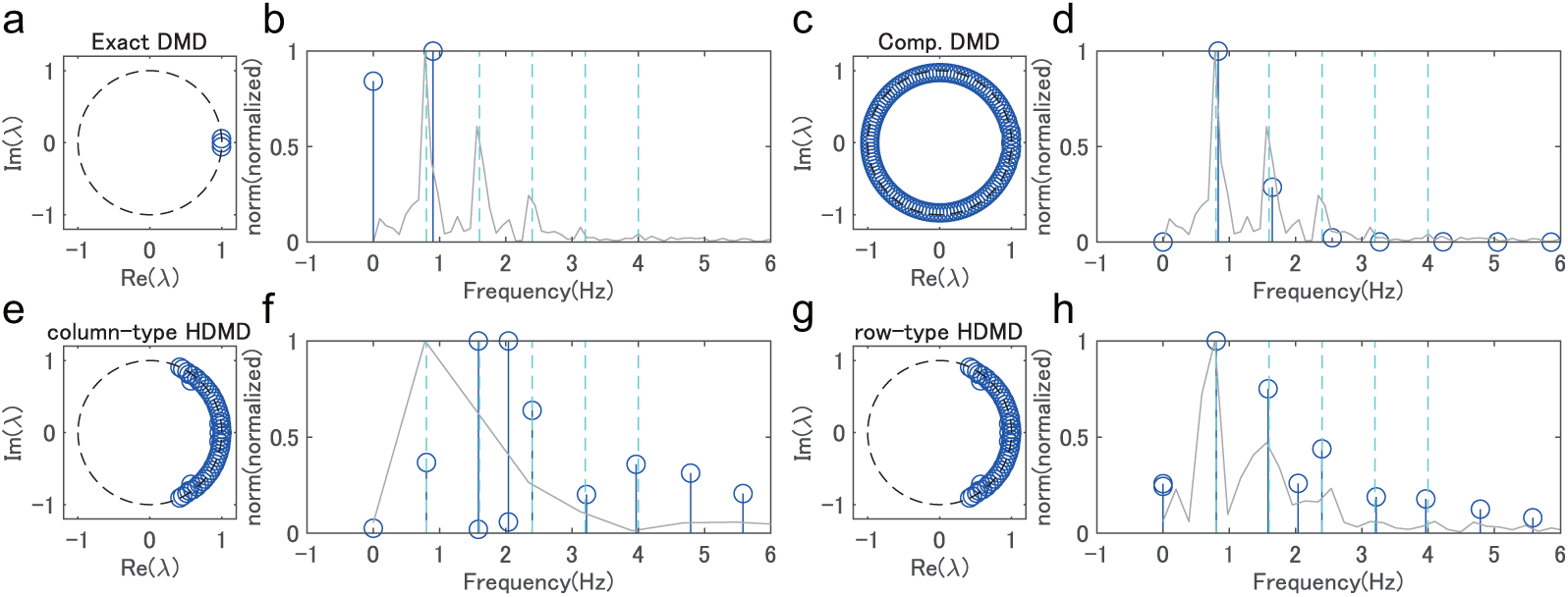
Representative examples of eigenvalues in four decomposition methods. Results of DMD eigenvalues in a complex plane (a,c,e,and g) and DMD and Fourier spectrum (b,d,f, and h) for exact DMD (a and b), companion-matrix DMD (c and d), column-(e and f) and row-type Hankel DMD (g and h) are shown. We used the same example of Fig 2. DMD and Fourier spectra are shown as blue and gray lines, respectively. Harmonic frequencies of the gait frequency are also shown as light blue dashed line.

Next, we visually and quantitatively evaluated the extraction of the harmonic frequencies in Fig 6. We show the averaged spectrum for all participants of the normalized spectrum of each sequence so that the maximal norm is 1. Since the gait frequencies during walking with the same speed were slightly different within and among participants, we normalized the frequency by dividing by the gait frequency. For all walking speeds, column- and row-type Hankel DMDs in Fig 6a and b visually obtained five harmonic frequencies (we denote the estimated five frequencies as *ω*_1_,…, *ω*_5_). However, companion-matrix DMD in Fig 6c shows (visually) only two harmonic frequencies. Note that we did not show the result of exact DMD because of the fewer extracted dynamics in Fig 5. Then, we quantitatively evaluated the difference between the obtained frequencies and the harmonic frequencies of the gait frequency after the normalization by dividing by the harmonic frequencies. These differences of companion-matrix DMD and column-type and row-type Hankel DMDs were 0.2655 ± 0.0427, 0.0214 ± 0.0100, and 0.0213 ± 0.0100, respectively. Since averaged Fourier spectra in Fig 6d were smoothed around the gait frequency (continuous spectra), we did not quantitatively evaluate the averaged Fourier spectra. Also, the results of the double pendulum simulation data shown in Fig S5 indicate that both types of Hankel DMDs extracted the two eigenfrequencies computed by the analytical solution. Moreover, in the walking simulation data shown in Fig S6, we obtained similar results to Fig 6.

**Figure 6:**
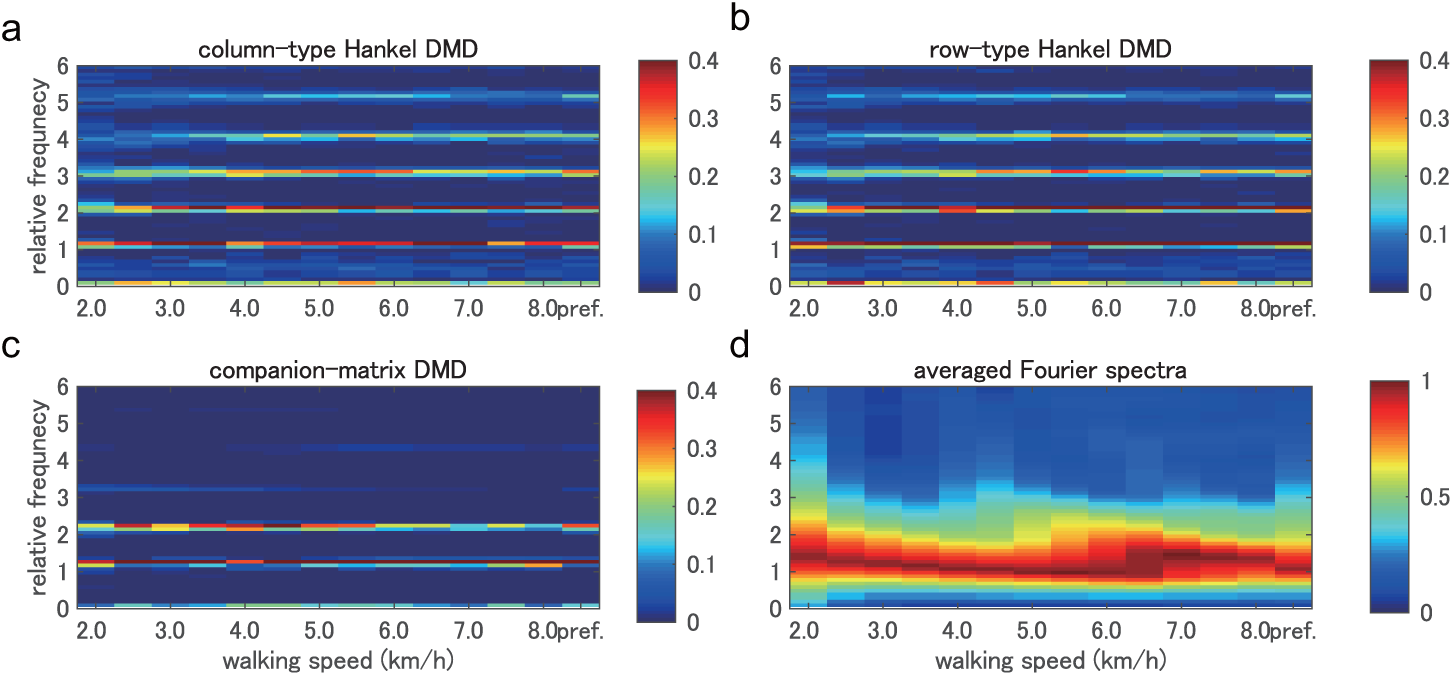
Extracted frequencies in four decomposition methods. Temporal frequency spectra in relative frequency domain to the gait frequency for (a) column-type and (b) row-type Hankel DMDs, (c) companion-matrix DMD, and (d) Fourier transformation averaged among all participants for all walking speeds are shown. Spectrum of each sequence was normalized so that the maximal norm is 1. For all walking speeds, column- and row-type Hankel DMDs visually obtained five harmonic frequencies. However, companion-matrix DMD visually shows only two harmonic frequencies. Averaged Fourier spectra were smoothed around the gait frequency.

### Koopman eigenfunctions

Next, in Fig 7, we visualized the speed-dependent time-evolving behavior of the phase of Koopman eigenfunction (in color) on the trajectory obtained by SVD-based method (i.e., in principal coordinates) to understand the dynamics on the conventional low-dimensional structure. Here we used row-type Hankel DMD (note that column-type Hankel DMD cannot extract the eigenfunction corresponding to the one-dimensional dynamics). The Koopman eigenfunctions 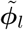 computed via row-type Hankel DMD are equivalent to the columns of the Hankel DMD modes corresponding to the focused dynamics (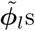 for *l* = 1,…, 5 correspond to *ω*_1_,…, *ω*_5_). We computed the argument of the complex-valued Koopman eigenfunction 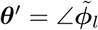 as shown in [36]. For clarity, we aligned the initial phases to near-zero values by the time-shift for all the eigenfunctions. Thus, all trajectories in Fig 7 for a participant rotate clock-wise from the black dots.

**Figure 7:**
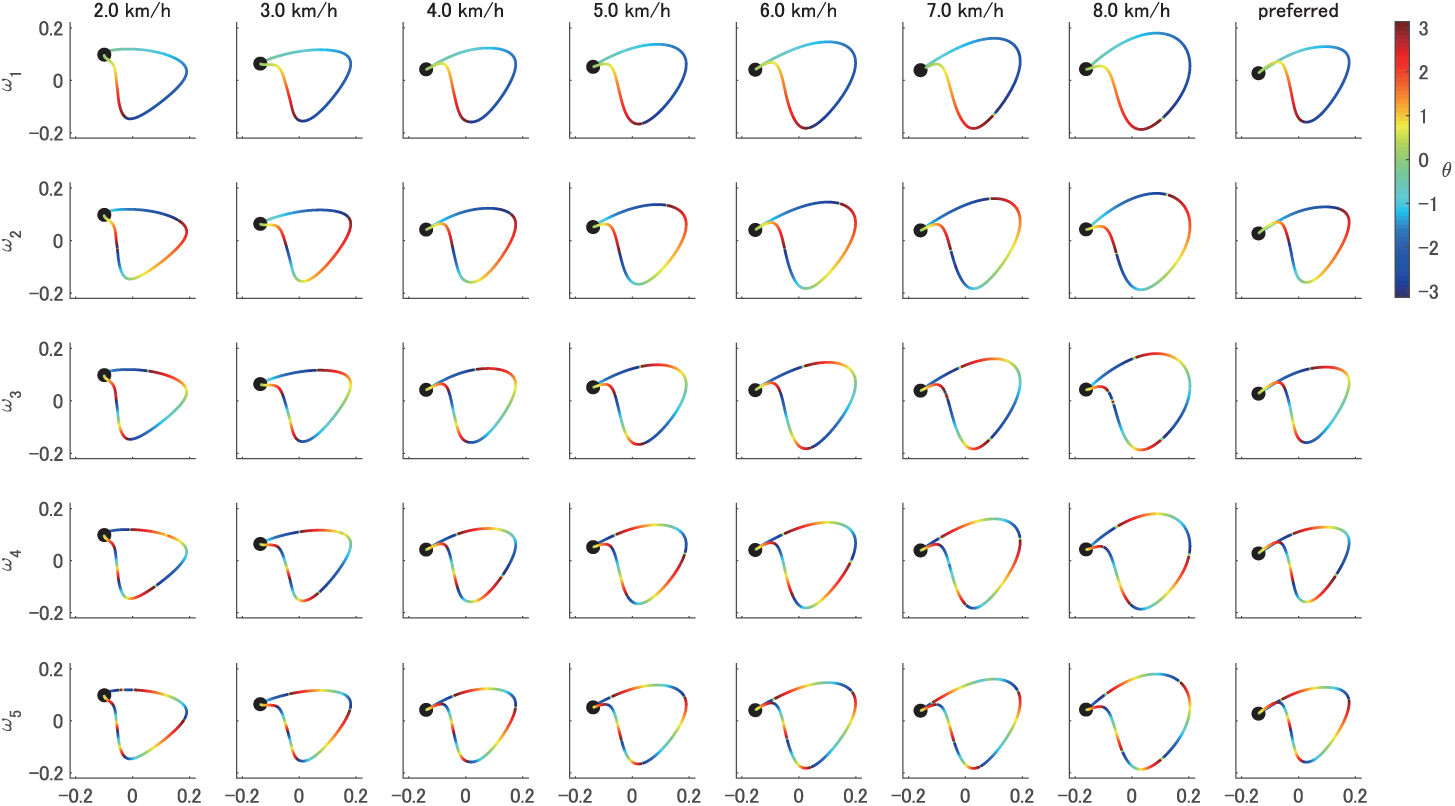
Phases of the Koopman eigenfunctions estimated by row-type Hankel DMD on the conventional coordinative structures. Phases computed as the argument of the Koopman eigenfunctions estimated by row-type Hankel DMD are shown for each walking speed and five harmonic frequencies of a participant. For clarity, we aligned the initial phases to near-zero values by the time-shift for the all eigenfunctions. All trajectories rotate clock-wise from the black dots. Although the shapes were similar among various velocities, the changes in the phase with walking speed (or gait cycles) were inhomogeneous. For example, the phases of *ω*_*l*_ visually advanced with the increase of the walking speed, such as the boundary between blue and red at the bottom in the plots of *ω*_1_ (but not the boundary between red and yellow).

Results show that, although the shapes in Fig 7 were similar among various velocities, the time-evolving behaviors of the phase with walking speed (or gait cycles) were heterogeneous. For example, the phases of *ω*_*l*_ visually advanced with the increase of the walking speed, such as the boundary between blue and red at the bottom in the plots of *ω*_1_ (but not the boundary between red and yellow). One of the possible reasons may be the decrease in the proportion of stance phase (the earlier phase in this case) and the increase in the proportion of swing phase (the later phase) as the walking speed increases [52]. This would be generated from the faster flexion/extension of the three joints as illustrated such as in [53, 54]. We also discuss this point further below.

Next, we quantitatively evaluated our results by comparing them to the eigenfunction obtained from another method called Fourier (or harmonic) average [38]. Fourier average requires a prior knowledge of the frequency (in this case, the accurate harmonic frequencies of the gait frequency) and the Koopman eigenfunction *ϕ*_*l*_ computed via Fourier average was the ideal oscillator without the growth/decay dynamics. We adjusted the initial phases of the Koopman eigenfunction 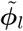 to those of *ϕ*_*l*_. The mean differences of two methods among all the time and sequences were small: for *ω*_1_,…, *ω*_5_ among all speeds and participants, they were 0.0281 ± 0.0117, 0.0120 ± 0.0059, 0.0068 ± 0.0033, 0.0021 ± 0.0015, and 0.0025 ± 0.0013, respectively. We also visualized the time-evolving behavior of the phase for the double pendulum and walking model simulation data in Fig S7.

## 3. Discussion

The objective of this study was to reveal dynamical properties of coodinative structures in biological periodic systems with unknown and redundant dynamics by a data-driven spectral analysis of dynamical systems (i.e., Hankel DMD). In this section, for each paragraph, we discuss the results of the extracted harmonics of gait frequencies, the comparison with the conventional coordinative structures, the extracted properties as dynamical systems, and the future perspectives. We then follow up with conclusions.

First, we extracted the harmonics of the gait frequency for various walking speeds in Fig 6. Especially, both column- and row-type Hankel DMDs extracted the distinct first to fifth harmonics compared with other DMD methods. In general, a periodic function that satisfies Dirichlet’s conditions can be expanded in the harmonic trigonometric functions with constant offset (e.g., [55]). Regarding (approximately) linear systems, for example, the double pendulum with a small initial condition in Fig S5, indicates only obvious frequency spectra. However, for nonlinear periodic systems, the strength of the spectra of the harmonics is not obvious, whereas the existence of the harmonics is obvious. Although the harmonics during gaits has been previously observed such as in ground reaction forces [56] and head movements [57], there is a possibility of measurement noise to generate the harmonics of the gait frequency [58]. Then, we compared the results of walking model simulation without the measurement noise, and confirmed the similar harmonics in Fig S6, suggesting that the harmonics would be generated from the interactions of multi-link and with environments (e.g., a ground and gravity) rather than the measurement noise. For such a redundant dynamical system, the equations of motion of the link model (described in Materials and Methods) become complicated as the model approximates a human. The data-driven method in this study to extract the dynamic properties of the coordinative structures would be beneficial for the understanding of the underlying dynamics.

Compared with the conventional coordinative structures, Hankel DMDs extracted dynamical properties behind the data. The conventional SVD-based method, which projects data on a low-dimensional subspace with orthogonal basis fitted by the data, showed better performance in terms of reconstruction error than DMDs but did not extract dynamic properties. These two approaches have different strengths and cannot be fairly compared. Instead, we can comprehensively understand the coordinative structures extracted by the two approaches. Then, we here visualized the relation between coordinative structures by SVD-based method (Fig 8a and b) and column-type Hankel DMD (Fig 8c-g). The variance along the third axis (PC3 in Fig 8b) in the space spanned by three principal directions using SVD indicates the deviation from the existing two-dimensional plane [10, 21, 22]. Also in the harmonic frequencies obtained by column-type Hankel DMD in Fig 8c-g, the deviation in the PC3 axis was smaller (on average among all harmonic frequencies, participants, and speeds, maximal absolute deviation was 0.0159 ± 0.0101) than that of SVD (0.1469 ± 0.0251). These results indicate that the reconstructed dynamics from Hankel DMD with the harmonic frequencies also lie in the conventional low-dimensional subspace. However, it should be noted that a similar spatial structure on the plane among various walking speeds did not mean the similar phase of the limit cycle, as shown in Fig 7. In other words, the conventional SVD-based method (e.g., [59, 28]) often normalized the time series to obtain consistent results, but this loses information on time or phase of the limit cycle. In general, at high harmonic frequencies, normalized time-series seems to be delayed when walking speed increases (but actually advanced because of the higher gait frequency or shorter gait cycle) as shown in Fig 9. Our approach using operator-theoretic spectral analysis can be useful because it can define the phase in terms of the underlying nonlinear dynamical systems.

**Figure 8:**
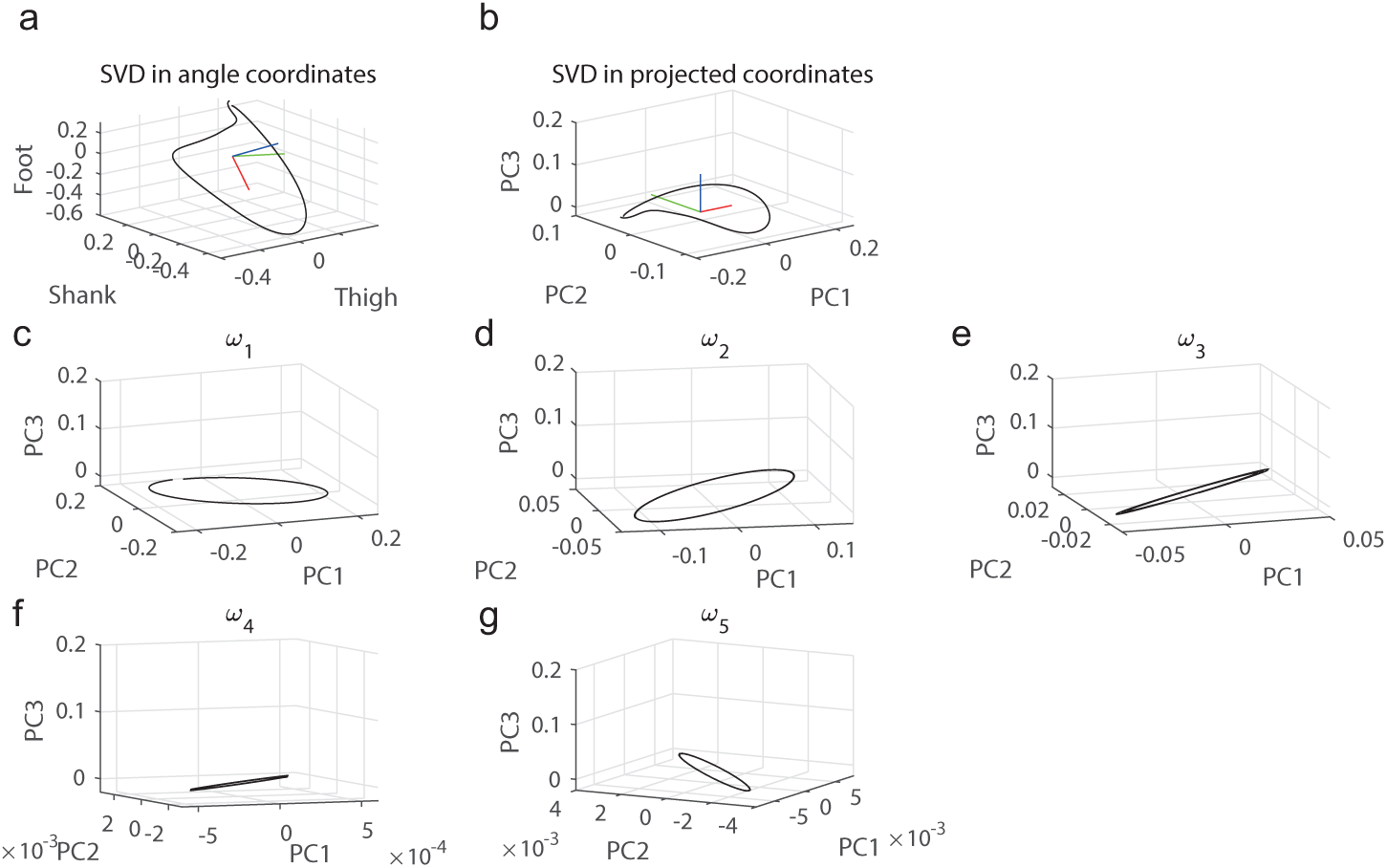
Relation to existing coordinative structure. (a) Trajectory in three angle coordinates with the project coordinates (red, green, and blue indicate the first, second, and third directions of principal components (PC1, PC2, and PC3) computed by existing SVD-based method. (b) Trajectory in projected coordinates by rotating (a). (c-g) Trajectories of five harmonic frequencies in the projected coordinates.

**Figure 9:**
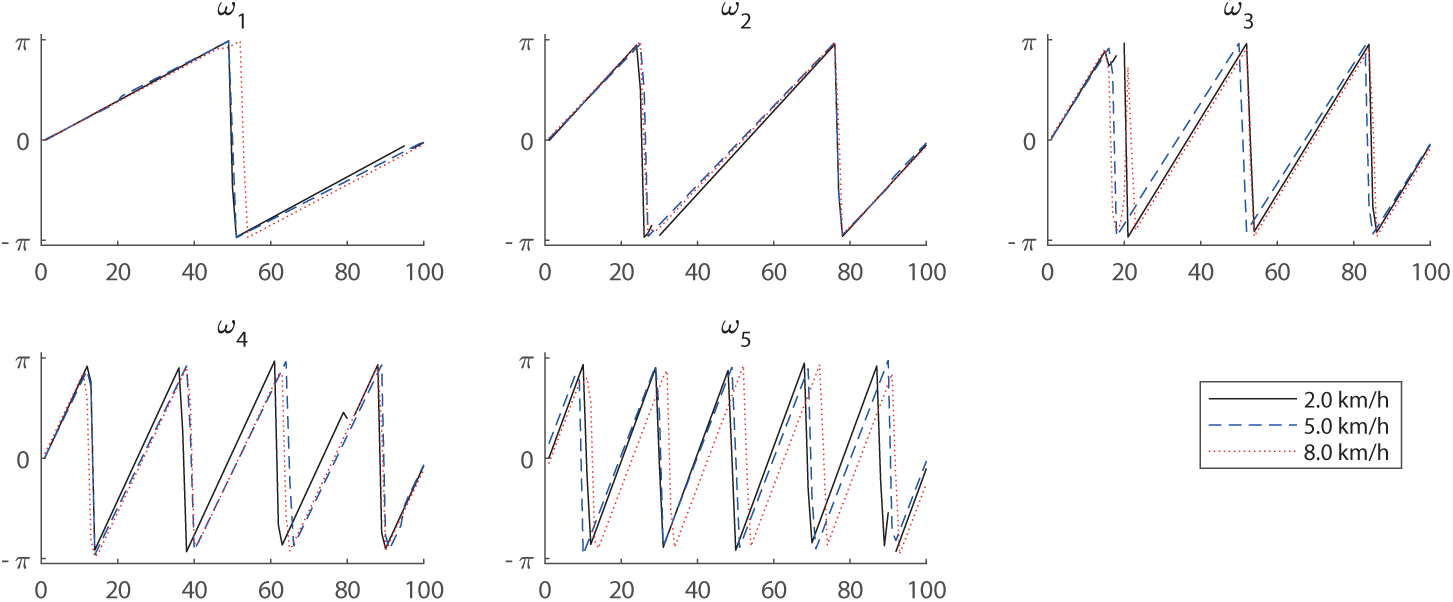
Normalized time-series of the phases of Koopman eigenfunctions. Phases computed by the argument of the Koopman eigenfunctions estimated by row-type Hankel DMD are shown for three walking speeds and five harmonic frequencies (a-e). Time series were normalized to 100 time stamps. For clarity, we aligned the initial phases to near-zero values by the time-shift for the all eigenfunctions (non-continuous sequences are the byproduct of the time-shift). The phases visually delayed (but actually advanced because of the higher gait frequency or shorter gait cycle) with the increase of the walking speed.

To investigate the dynamical properties of the underlying dynamical systems such as a limit cycle, our approach has advantages because it can estimate Koopman eigenvalues (frequency or decay/growth rate), eigenfunctions (projection from data space to latent state space), and DMD modes (spatial coefficients). For understanding biological redundant systems as dynamical systems, it should be noted that the governing equations including neuro-musculo-skeletal dynamics are often partly unknown, which would be different from most of the known physical systems. In this case, as discussed above, a data-driven approach such as used in this study would be effective. Other data-driven approaches may be also effective such as curve fitting using the dictionary of the basis functions (e.g., [60]), and actually, researchers have applied a sinusoidal curve fitting to the intersegmental coordination during locomotion [61]. Meanwhile, our approach is the data-driven operator-theoretic spectral analysis of nonlinear dynamical systems without prior explicit knowledge. We adopted this approach with the assumption that there are underlying dynamical systems or limit cycles behind the segmental angles during locomotion. In limit cycles, phase description has been developed for locomotion such as using local joint angles [47, 48, 49, 13] or right heel-contact cycle as global description [50]. Compared with these studies, we extracted global phases at the gait frequency and at additional harmonic frequencies in a data-driven manner. However, although we can interpret that the double harmonic frequency may relate to a one-step movement (because one gait cycle includes two steps) and higher harmonic frequencies may relate to the impact and/or absorption of the foot contact, we currently did not provide strict mechanical interpretations.

Further studies would be needed at least from three perspectives. One is the application to a control problem using forward simulations. The global entrainment among the neuro-musculo-skeletal system and the environment during locomotion should be further investigated by neuro-musculo-skeletal simulations with neural inputs such as central pattern generator [4] and reflex action [8] by feedback from environments. These will contribute to the understanding of the above mechanical interpretation of the harmonic frequencies. Second is related to our assumption that the walking dynamics is on a limit cycle. Actually, walking is sometimes not an ideal cyclic motion; thus we need a method adapting to rhythm shift and perturbation (observed as a phase reset in [47, 50, 48] and a phase locking in [49]). Our approach can theoretically describe an asymptotic phase in asymptotically stable dynamical systems [38, 36]; therefore, it will be possible to apply it to the perturbed or more complicated locomotion [62, 63]. Third is the variation within/among participants. Although this study basically investigated the difference among walking speeds, this dynamics directly related to the gait (and harmonic) frequency. Comparison with other types of locomotion, or other specific participant groups will contribute to the understanding of the redundant and unknown human motor systems as dynamical systems.

In conclusion, we applied data-driven and operator-theoretic spectral analysis to intersegmental angles during human locomotion, which can extract coordinative structures based on dynamic properties and obtain a phase reduction model. We adopted Hankel DMD, which theoretically yields the eigenvalues and eigenfunctions of composition (Koopman) operator by augmenting finite data. First, we extracted characteristic harmonic frequencies from intersegmental angles during human walking at various speeds. Second, we discovered the speed-dependent time-evolving behaviors of the phase on the conventional low-dimensional coordinative structure by estimating the eigenfunctions. We also verified our approach using the double pendulum and walking model simulation data. Our approach contributes to the understanding of unknown and redundant periodic phenomena of living organisms from the perspective of nonlinear dynamical systems.

## 4. Materials and Methods

### Participants

Ten healthy men (age: 23.3 ± 0.9; height: 171.1 ± 3.44 cm; weight: 64.1 ± 0.63 kg) participated in this study. The participants provided written informed consent to participate in the study after receiving a detailed explanation of the purpose, potential benefits, and risks associated with participation. The experimental procedures were conducted in accordance with the Declaration of Helsinki and were approved by the Local Ethics Committee of the Graduate School of Human and Environmental Studies, Kyoto University (Approval number 26-H-22).

### Experimental setup and data preprocessing

Participants walked on a treadmill (Adventure 3 PLUS, Horizon, Johnson Health Tech Japan Co., Tokyo, Japan) at 14 different controlled speeds (2.0, 2.5, 3.0, 3.5, 4.0, 4.5, 5.0, 5.5, 6.0, 6.5, 7.0, 7.5, and 8.0 km/h and a preferred walking speed of 4.3 ± 0.63 km/h) that were administered in a random order over the span of 50 gait cycles. To determine the preferred walking speed for each individual participant, we modulated the treadmill speed without showing the walking speed to the participants. The preferred walking speeds were determined at the moment when the participants felt comfortable.

Kinematic data were recorded using a 3D optical motion capture system with 12 cameras operating at 100 Hz (Optotrak, Northern Digital Inc., Waterloo, Ontario, Canada). This system captured three-dimensional coordinates of reflective markers that were attached to the anatomical landmarks on the participants. The reflective markers that were attached to the participants were positioned at the top, right and left sides of their heads, as well as on the acromions, elbows, wrists, anterior superior iliacspine, posterior superior iliac spine, greater trochanters, medial and lateral epicondyles, medial and lateral malleolus, heels and toes. The measured reflective marker data were low-pass filtered at 8 Hz, which contains the frequency band in the following analyses. We defined one gait cycle as the time between the initial right heel contact and the next right heel contact in a previous method [64]. We primarily used one gait cycle for the analysis (for details, see Hankel DMD subsection). Similarly, we defined gait frequency as the reciprocal number of a gait cycle. We defined six elevation angles (Fig 1a and b: right and left feet, shanks, and thighs). Our selection of elevation angles was based on the assumption that changes in elevation angles are more stereotypical than relative angles [10, 51]. To prepare to extract coordinative structures, we computed mean postures by averaging the elevation angles over a gait cycle and subtracting them from elevation angles for each segment angle [59, 28]. For the subsequent analysis, we primarily used three angles (right feet, shank, and thigh) to compare the conventional coordinative structures (e.g., [10, 51]), but when we simulated the following five-link biped model, we used four angles (right and left shanks and thighs). We divided all data into validation and test datasets including both 10 sequences. We used the validation dataset in the following procedure of determining the parameters of Hankel DMD. Thereafter, we performed all of the remaining analysis using the test dataset.

### Conventional decomposition method based on SVD

The conventional intersegmental coordination (also called kinematic synergies) was extracted from the pre-processed elevation angle time-series matrix **Θ** *∈* ℝ^*d×τ*^ (*d* = 3, 4 and *τ* is the time length) by SVD [59, 28]. We call it SVD-based method in this study. The conventional two-dimensional coordinative structure (i.e., a plane) in three-dimensional angle space [10, 21, 22] can also be extracted by the same procedure. By applying SVD to the processed elevation angle matrix, the matrix was decomposed into the intersegmental coordination ***z***_*j*_ and the temporal coordination (*λ*_*j*_***ν***_*j*_)^T^ such that

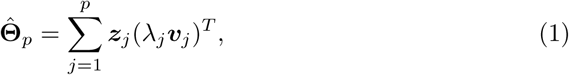

where *p* is the number of intersegmental coordination (0 ≥ *p* ≥ *d*). Intersegmental coordination ***z***_*j*_ represents the principal groups of the *j*th simultaneously active segmental group, and temporal coordination *λ*_*j*_***ν***_*j*_ indicates the activation patterns of *j*th intersegmental coordination. To evaluate the dimension of the reduction or determine the number of intersegmental coordination, researchers used the cumulative contribution ratio using eigenvalues (e.g., [59, 28]) because of the orthogonal decomposition. However, in this study, to compare with non-orthogonal decomposition methods such as DMD, we adopted VAF which is commonly used in non-orthogonal dimensionality reduction such as non-negative matrix factorization [65] in muscle activity. VAF is defined as the square error of the reconstructed data and the original data such that 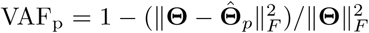 where ||·||_*F*_ is the Frobenius norm. There are some procedures to determine the number of dimensions of the structure based on such as VAF, but for comparison with DMDs, we set *p* = 2 for the actual human locomotion and pendulum simulation data and *p* = 3 for the walking model simulation data. Moreover, for comparison with DMDs, since the augmented data dimension is variable in Hankel DMD, the absolute reconstruction error is defined as 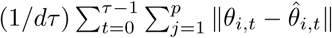, where *θ*_*i,t*_ and 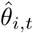 are scalar components of **Θ** and 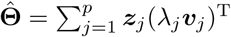 at dimension *i* and time *t*, respectively.

### Koopman spectral analysis and variants of DMDs

Here, we first briefly review Koopman spectral analysis, which is the underlying theory for DMD, and then describe the basic DMD procedure. First, we consider a nonlinear dynamical system: ***x***_*t*+1_ = ***f*** (***x***_*t*_), where ***x***_*t*_ is the state vector in the state space ℳ *⊂* ℝ^*p*^ with time index *t ∈* 𝕋: = {0} *∪* ℕ. *The Koopman operator*, which we denote by 𝒦, is a linear operator acting on a scalar observable function *g*: ℳ *→* ℂ defined by

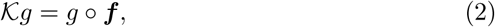

where *g* ∘ ***f*** denotes the composition of *g* with ***f*** [29]. That is, it maps *g* to the new function *g* ∘ ***f***. We assume that *𝒦* has only discrete spectra. Then, it generally performs an eigenvalue decomposition: *𝒦ϕ*_*j*_(***x***) = *λ*_*j*_*ϕ*_*j*_(***x***), where *λ*_*j*_ *∈* ℂ is the *j* -th eigenvalue (called *the Koopman eigenvalue*) and *ϕ*_*j*_ is the corresponding eigenfunction (called *the Koopman eigenfunction*). We denote the concatenation of scalar functions as ***g***: = [*g*_1_,…, *g*_*d*_]^T^. If each *g*_*i*_ lies within the space spanned by the eigenfunction *ϕ*_*j*_, we can expand the vector-valued ***g*** in terms of these eigenfunctions as 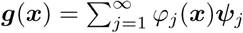, where *ψ*_*j*_ is a set of vector coefficients called *the Koopman modes*. Through the iterative applications of 𝒦, the following equation is obtained:

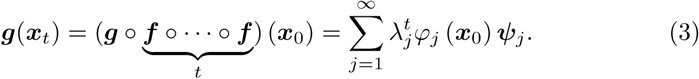

Therefore, *λ*_*j*_ characterizes the time evolution of the corresponding Koopman mode *ψ*_*j*_, i.e., the phase of *λ*_*j*_ determines its frequency and the magnitude determines the growth rate of its dynamics.

Among several possible methods to compute the above modal decomposition from data, DMD [32, 33] is the most popular algorithm, which estimates an approximation of the decomposition in Eq. 3. Consider a finite-length observation sequence ***y***_0_, ***y***_1_,…, ***y***_*τ*_ (*∈*ℂ^*d*^), where ***y***: = ***g***(***x***_*t*_). Let ***X*** = [***y***_0_, ***y***_1_,…, ***y***_*τ*– 1_] and ***Y*** = [***y***_1_, ***y***_2_,…, ***y***_*τ*_]. Then, DMD basically approximates it by calculating the eigendecomposition of matrix ***F*** = ***Y X***^*†*^, where ***X***^*†*^ is the pseudo-inverse of ***X***. The matrix ***F*** may be intractable to analyze directly when the dimension is large. Therefore, in the popular implementation of DMD called exact DMD [66], a rank-reduced representation 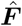 based on SVD is applied. That is, ***X*** *≈* ***U* ∑ *V*** ^*\**^ and 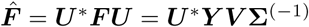, where ^*\**^ is the conjugate transpose. Thereafter, we perform eigendecomposition of 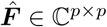 to obtain the set of the eigenvalues *λ*_*j*_ and eigenvectors ***w***_*j*_. Then, we estimate the Koopman modes in Eq. 3: 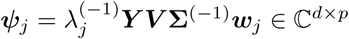, which is called *DMD modes*.

In this study, we consider real-valued input matrices ***X***, ***Y*** *∈* ℝ^*d×τ*^, whereas DMD can be applied to complex-valued matrices. Even if we use the real-valued input matrices, since 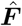 is usually a non-Hermitian matrix (in a real-valued case, an asymmetric matrix), eigenvalues and eigenvalues (then, DMD modes) are often complex. Therefore, we used the absolute value of *ψ*_*i,j*_, which is a component of *ψ*_*j*_ for *i* = 1,…, *d*, for representing the spatial weight of the intersegmental coordinative structures. Eigenvalue *λ*_*j*_ is transformed into temporal frequency *ω*_*j*_ such that *ω*_*j*_ = ln(*λ*_*j*_)*/*2*π*Δ*t*, where Δ*t* is a time interval of the discrete time system. Time dynamics of *j*th mode, which corresponds to the temporal coordination in SVD-based method, is defined as exp(*ω*_*j*_*t/*2*π*)***b***_0_, where 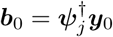 [67] and 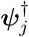 is the pseudoinverse of *ψ*_*j*_. For comparison with SVD-based method and Hankel DMD, the absolute reconstruction error is defined as 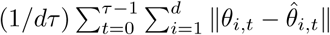, where *θ*_*i,t*_ and 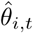 are scalar components of **Θ** and 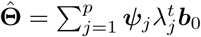 at dimension *i* and time *t*, respectively. In addition, we defined VAF in the same way as the case of SVD-based method.

We also examined another popular DMD implementation called companion-matrix DMD (the name is based on [36]). Companion-matrix DMD, originally developed in [32], is a more straightforward formulation than exact DMD, but sometimes numerically unstable. Algorithmically, the number of DMD eigenvalues *p* in exact DMD is limited to the data dimension *d* if *d ≪ τ* such as in this case, whereas that in companion-matrix DMD is limited to the time length *τ* in this case (note that DMD algorithms basically compute pairs of complex conjugate eigenvalues or dynamic modes). This property in exact DMD suffers from the problem when we use some biological data including more dynamic modes than the data dimension such as multi-link motion in this study. Theoretically, each observable function *g*_*i*_ should lie within the space spanned by the Koopman eigenfunction *ϕ*_*j*_, i.e., the data should be rich enough to approximate the eigenfunctions. However, in basic DMD algorithms naively using the obtained data such as both exact and companion-matrix DMD, the above assumption is not satisfied such as when the data dimension is too small to approximate the eigenfunctions.

Then, there are several algorithmic variants to overcome such problem of the original DMD such as the use of a dictionary of basis functions (e.g., [68]), a formulation in a reproducing kernel Hilbert space [69]. These methods work well when appropriate basis functions or kernels are prepared; however, it is not always possible to prepare such functions if we have no prior knowledge of the underlying dynamics. Another variant has been recently developed using nonlinear transformation in a neural network framework (e.g., [70]), which does not need prior knowledge about the observable function. However, in this study, we assume that the dynamical system behind the angle series data during walking is a limit cycle, which is a relatively tractable class in dynamical systems. Thus, we adopt a more straightforward approach without nonlinear transformation according to the previous work including a limit cycle [71, 36] or unknown dynamics such as [35, 34]. Hankel DMD [36] is also a variant of DMD applied to delay embedding matrix, which can theoretically yield Koopman eigenvalues and eigenfunctions. The Koopman eigenfunction provides linearly evolving coordinates in the underlying state space and can describe the phase in (asymptotically) stable dynamical systems [38, 36]. Therefore, we adopted Hankel DMD, as described in the next subsection.

### Hankel DMD

Here, we introduce Hankel DMD and explain the procedure in this study. Hankel DMD uses the Hankel matrices of data in the forms of

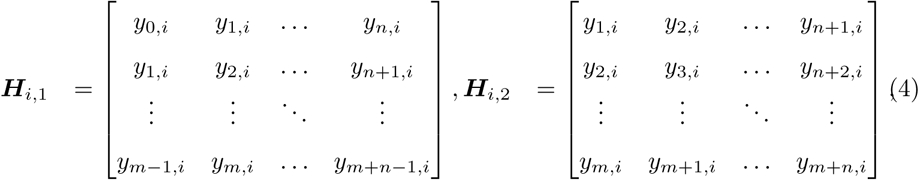

for *i* = 1, 2,…, *d*, where *y*_*t,i*_ is a component of vector ***y***_*t*_ *∈* ℂ^*d*^ for *t* = 0,…, *τ*. Although in [36] the scaling factors were computed for normalizing the norms of the observables, the segmental angles in this study were not extremely large or small for each other (Fig 1b); thus we did not use the scaling factors. Next, we form the concatenated matrices:

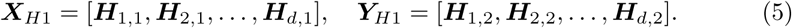

Then, we compute the truncated 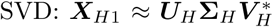 and obtain 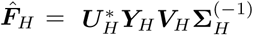 as the exact DMD procedure. Thereafter, we perform eigende-composition of 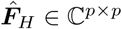 to obtain the set of eigenvalues *λ*_*j*_ and eigenvectors ***w***_*j*_ for *j* = 1,…, *r*. A set of *λ*_*j*_s approximates the Koopman eigenvalues. We call it row-type Hankel DMD. Conventional Hankel DMD modes are given by: 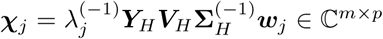. From [36], the conventional Hankel DMD mode ***χ***_*j*_ converges to the Koopman eigenfunctions 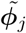, which are the columns of ***χ***_*j*_ corresponding to the focused dynamics, if *m* is sufficiently large. We compute this row-type Hankel DMD for estimating Koopman eigenfunctions corresponding to one-dimensional dynamics. For extracting the coordinative structure among observables (e.g., segmental angles), we have a problem that the conventional Hankel DMD modes ***χ***_*j*_ *∈* ℂ^*m×p*^ cannot directly compute the coordinative structure between elements of a vector-valued observable (e.g., segmental angles). Then, we need to define row-type Hankel DMD modes regarding original observables (no duplication by delay embedding). Note that in general, we have several choices to define DMD modes (e.g., reviewed by [36]) in which the mode and the estimated value of Koopman eigenfunctions (here we denoted ***ϕ***_*j*_ *∈* ℂ^*m*^ and 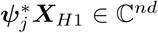, respectively) satisfy 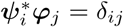 is the Kronecker’s delta). Then, since conventional Hankel DMD mode ***χ***_*j*_ is here used as eigenfunction 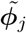, we defined the complementary quantities, i.e., the row-type Hankel DMD modes, from the vector corresponding to the conventional eigenfunction value 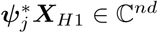 satisfying 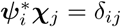. Finally, we selected the row-type Hankel DMD mode regarding original observables (without duplication) from 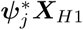 by picking up the values corresponding to the initial value for each dimension.

The coordinative structure, can be computed in a more straightforward way as follows. We define the concatenated matrices as input matrix of Hankel DMD such that

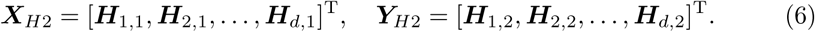

We call it column-type Hankel DMD, which is similar to the method such as used in [35] with delay-coordinated matrices. For the above procedures, we can compute column-type Hankel DMD modes and eigenvalues in the similar procedure as exact DMD. However, column-type Hankel DMD modes duplicate by the delay embedding; thus, when we compare with basic DMDs and SVD-based method, we picked up the Hankel DMD modes without the delay for each dimension. Note that it is difficult for column-type Hankel DMD to estimate a one-dimensional eigenfunction for each dynamics from the viewpoint of phase model. VAF and reconstruction error were also computed in the same procedure above. We set the time length *n* to one gait cycle *T* frames, in order to compare the result of the previous works in walking intersegmental coordination such as [51, 59]. In [36], it was mentioned that a few hundred samples would be enough to determine the Koopman eigenvalues of periodic systems with high accuracy; thus, we fixed *n* = *T* and selected other parameters *p* and *m* as the following procedure.

For both types of Hankel DMDs, we need to determine the dimension *p* of truncated SVD and the dimension *m* of delay embedding. In this study, we determined *p* and *m* by considering both the convergence of Hankel DMDs as spectral analysis of a Koopman operator and the avoidance of fitting to high-frequency dynamics. It should be noted that although we performed cross-validation widely used such as in statistics and machine learning area, it conflicted with the above criteria, which is shown as an independent numerical experiment in Appendix A. Again, we divided data into validation and test datasets, and used only the validation datasets for the determination of *p* and *m*.

First, from the viewpoint of the convergence [36], the dimension *p* of truncated SVD theoretically converges to the dimension of the Koopman invariant subspace *k* if *m → ∞*. Practically, the previous work [36] used the hard threshold of SVD (1e-10). Another study in a similar algorithm [72] used the optimal hard threshold of SVD [73] when the noise level is unknown. Note that, obviously, the threshold of SVD is directly related to the dimension *p* of truncated SVD. For the dimension *m* of delay embedding, theoretically, sufficient *m* can approximate DMD modes to Koopman eigenfunctions (the necessary condition is obviously *m > k* + 1) [36]. However, the effect of *p* (or the SVD threshold) and *m* on the convergence is not unclear in the actual locomotion data. Therefore, we examined various *p*s and *m*s to find the sufficient (not optimal) *p* and *m* using the validation dataset as a guide for the biological studies. Here, we used the convergence of the reconstruction error (defined above) for investigating the convergence of the estimation error of Koopman eigenvalues and modes.

Second, from the viewpoint of stably obtaining the desirable DMD results, we empirically know that Hankel DMDs with too large *m* and *p* generate eigenvalues with too high frequencies. This may cause undesirable fitting to too high-frequency dynamics (i.e., DMD compute too large coefficients of the high-frequency dynamics) which should not be realistically considered. Thus, we visually selected certain *m* and *p* as small as possible while satisfying the condition in which the error converges.

For auxiliary information, it should be mentioned that there can be other approaches to determine *m*. In dynamical systems, determining delay embedding dimension has been discussed (e.g., reviewed by [74]) such as using false nearest neighbors [75] as used in the analysis of joint angles during locomotion [76]. However, these approaches can be directly applied to dynamical systems but may not be guaranteed to be suitable for DMDs. Among the approaches for nonlinear dynamical systems to fit the data to basis functions (e.g., [60]), the study by [77] used Akaike information criteria [78] after sparse identification. However, again, we did not choose *m* and *p* from the perspective of minimizing the generalization error; thus, we did not use any information criterion.

### Double pendulum simulation

As a validation of our approach using a physical phenomenon with a multi-link structure which obeys known ordinary differential equations with analytically-obtained frequency modes, we used a double pendulum model. Generally, the motion of a double pendulum is chaotic, but this study used a small initial condition and then the equation can be linearized and obtained two eigenfrequencies. For simplicity, we considered a double pendulum consisting of two pendulums with the same length *l* and weight *m* attached end to end, as shown in Fig 1c. Parameters *θ*_1_ and *θ*_2_ are also in Fig 1c (we set *m* = 1 kg and *l* = 1 m). The governing equations are as follows:

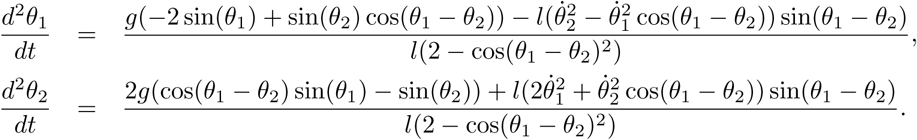

For small initial conditions (we set *θ*_1_ = *θ*_2_ = *π/*8 and 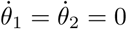), we obtain approximated linear systems

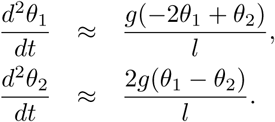

By solving the secular equation, we analytically obtain two eigenfrequencies 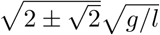. We verified our approach using the simple model with explicit governing equation and frequency modes. In the simulation, the time step and the duration were set to 1*/*20 s and 500 s in total. We set the analyzed interval to 160 time points based on Fig 1d (here, we did not use the information of the true frequencies). Similarly to the actual locomotion data, we divided all data into a validation dataset (10 sequences) for determining the parameters of Hankel DMDs and a test dataset (10 sequences) for the remaining analyses.

### Walking model and simulation

Next, to understand the equations of motion in locomotion and to verify our approach using a model simulation data with explicit governing equation (without noise) but with unknown frequency modes, we describe a simple human multi-link model. First, we describe a general overview of the dynamics, and indicate that the gait dynamics can be partly approximated to be a nonlinear function of the segmental angles ***θ***. The dynamics of a multi-link model are derived in the following Lagrangian equations:

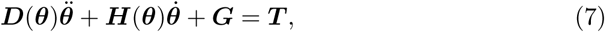

where ***D***(***θ***) is a positive definite and symmetric inertia matrix, ***H***(***θ***) is the matrix of a centrifugal and Coriolis terms, 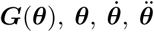 are a gravity term, generalized coordinates, velocities and accelerations, respectively (more details can be found such as in [4]). ***T*** is a generalized torque term, that has been complicatedly modeled in general but precisely unknown in actual humans.

Next, Eq. 7 can be transformed into the following form:

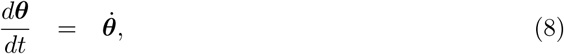

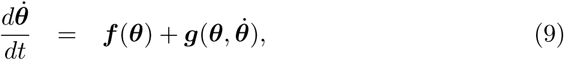

where ***f*** (***θ***) = ***D***(***θ***)^−1^(***T*** – ***G*(*θ*)**) and 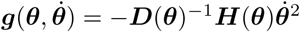. The results of previous work [20] in the numerical simulation showed that 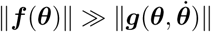. In other words, ***f*** (***θ***) represents the absolute domination in the dynamics of the biped model. Therefore, gait dynamics can be approximately represented by function ***f*** (***θ***) = ***D***(***θ***)^−1^(***T*** – ***G*(*θ*)**) along the phase portrait of ***θ***. Hence, segmental angles are selected as the kinematic parameters, because the gait dynamics can be partly approximated to be a nonlinear function of the segmental angles, except for the torque term.

Next, as a walking model simulation for comparison with the actual walking data, we used a well-known five-link neural oscillator control model [4] for simplicity (again, here we used four angles: left and right thigh and shank angles). We used the cited body parameters and motion equations and set the constant input as 6. In the simulation, the time step and the duration were set to 10^−6^ s and 120 s in total, but thereafter we employed the down-sampling into 100 Hz for the computation of DMDs. We set the analyzed interval to a gait cycle similarly to the actual human locomotion. We also divided all data into validation datasets (10 sequences) for determining the parameters of Hankel DMD and test datasets (10 sequences) for the remaining analyses.

## 5. Acknowledgments

This work was supported by JSPS KAKENHI (Grant Numbers 18K18116 and 18H03287); JSPS Research Fellow (Grant Number 16J07348); the Japanese Council for Science, Technology and Innovation (CSTI); and the Cross-ministerial Strategic Innovation Promotion Program (SIP Project ID 14533567 Funding agency: Bio-oriented Technology Research Advancement Institution, NARO).

This work was supported by JSPS KAKENHI Grant Numbers 16K12995, 16H01548, and 18H03287.

## Supplementary Materials

### Appendix A. Cross-validation results of the dimensions of Hankel DMD

From the perspective of minimizing generalization errors, as an independent experiment of the main text, we also performed cross-validation to select *p* and *m* minimizing the estimation error. Note that the Hankel DMD procedure does not estimate (or train) any parameter. Then, again, we divided data into validation and test datasets, and used the validation datasets for the leave-one-out cross-validation (LOOCV). Since *p* depends on *m* (column-type Hankel DMD: *p* ≥ min(*md,n*) and row-type: *p* ≥ min(*m,n*), we first determined *m* and then *p* to minimize the estimation error. However, we did not use the criteria to determine *m* and *p*, because too large *m* and *p* can decrease the generalization errors but may cause the undesirable fitting to the too high frequencies.

Next, we show LOOCV results for various *m*s and *p*s using the box-plots. In Appendix 1 Figure 1a and c, *m* in both Hankel DMDs decreased with the walking speed, whereas *p* in both Hankel DMDs in Appendix 1 Figure 1 b and d Figs did not depend on the speed. These indicate that the selected *m*s in LOOCV seemed to be related to the above *m* = *T* and the selected *p*s in LOOCV were larger than the above *p* = 50. The average LOOCV error for all velocities and participants was 0.0012 ± 0.0012 and 0.0015 ± 0.0007 in column- and row-type Hankel DMDs, respectively. For comparison with the above *m* and *p*, both average errors converged, but in Appendix 1 Figure 1 b and d, some *p*s were too high and we considered that it may cause the undesirable fitting to the high-frequency dynamics. Therefore, we did not select *m* and *p* using cross-validation.

## Supplementary Figures

**Figure S1:**
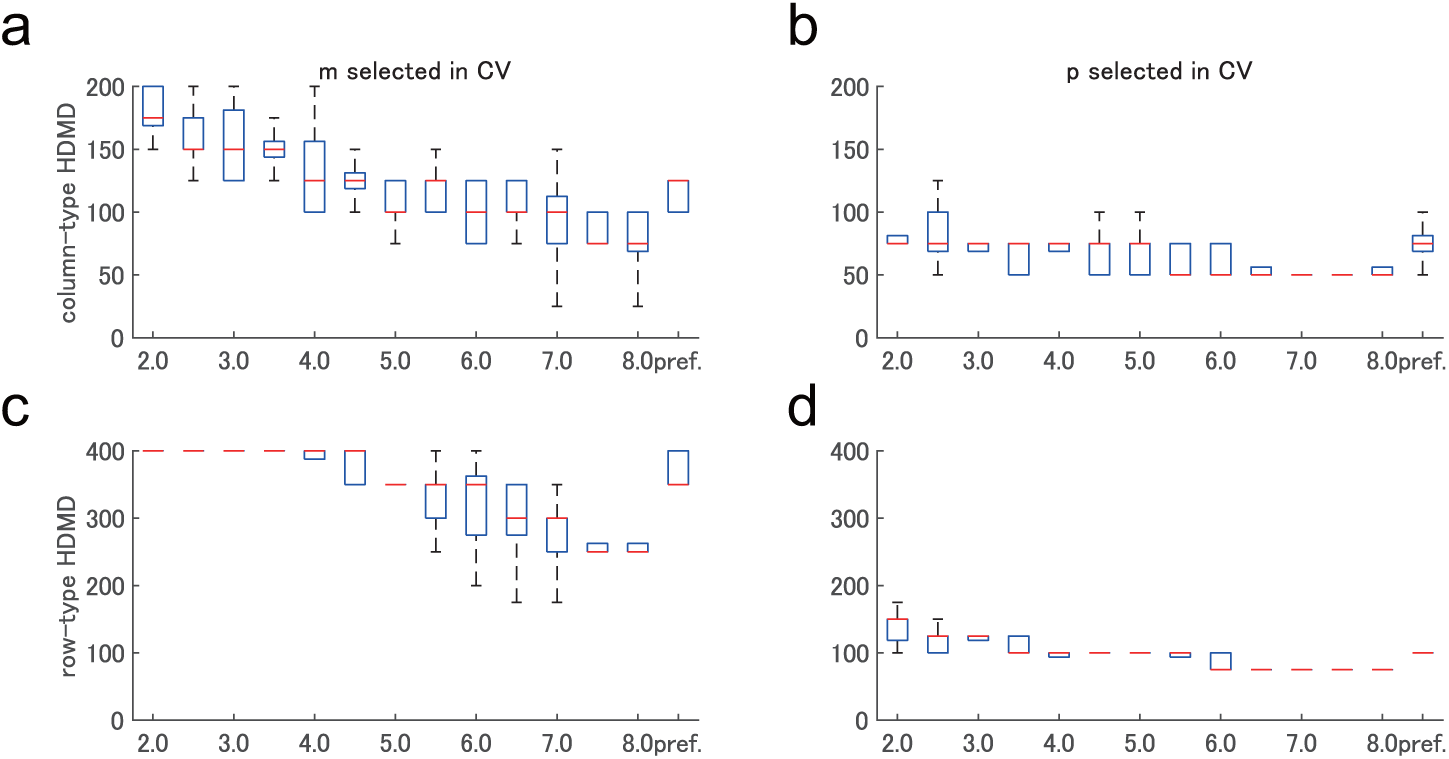
Selected dimensions *m* and *p* in cross-validation. Boxplots of selected dimensions *m* (a and c) and *p* (b and d) for column-type (a and b) and row-type (c and d) Hankel DMDs during various walking velocities are shown. The central mark (red) indicates the median, and the bottom and top edges of the box indicate the 25th and 75th percentiles, respectively. The whiskers extend to the most extreme data points.

**Figure S2:**
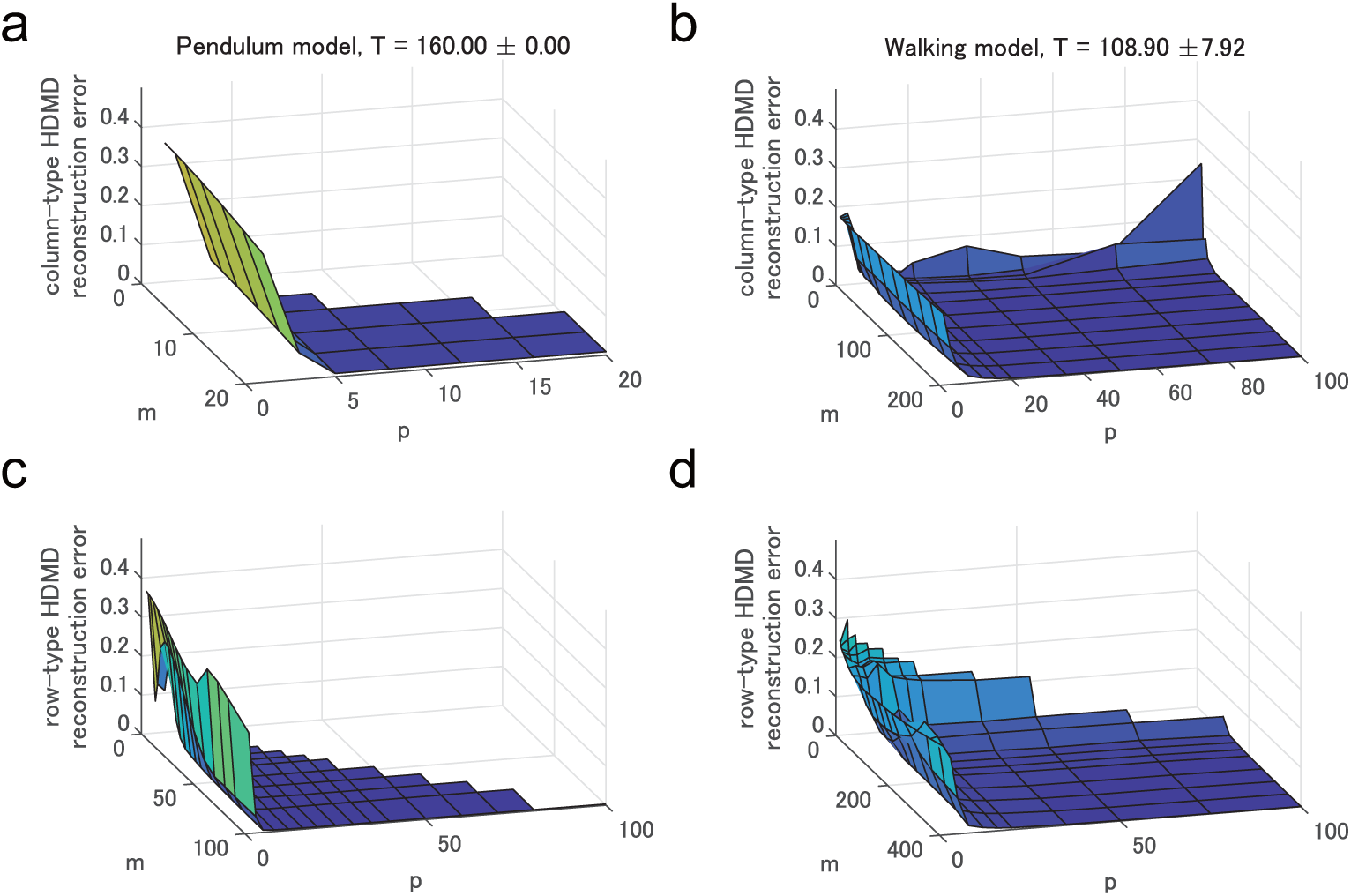
Convergence of Hankel DMDs for two simulation data. Examples of the reconstruction error for various *m* and *p* for column-type (a and b) and row-type (c and d) Hankel DMDs for the simulation data of the pendulum model and walking model are shown. Similarly to the human walking data, both column- and row-type Hankel DMDs with larger *m* and *p* converged to a certain error. We selected *p* = 20 and *m* = 20, 100 for column- and row-type Hankel DMDs for the pendulum model (error in column-type: 0.0090 ± 0.0047, error in row-type: 0.0031 ± 0.0004) and *p* = 50 and *m* = *T*, 2*T* for the walking model (error in column-type: 0.0021 ± 0.0013, error in row-type: 0.0073 ± 0.0102). For the pendulum model, we set *m* to 20 and 100 in the column-type and row-type because of the rank deficiency and the computation of the Koopman eigenfunction, respectively. The other criteria were the same as those described in the main text. The analyzed interval (*n*) is shown in (a and b) top.

**Figure S3:**
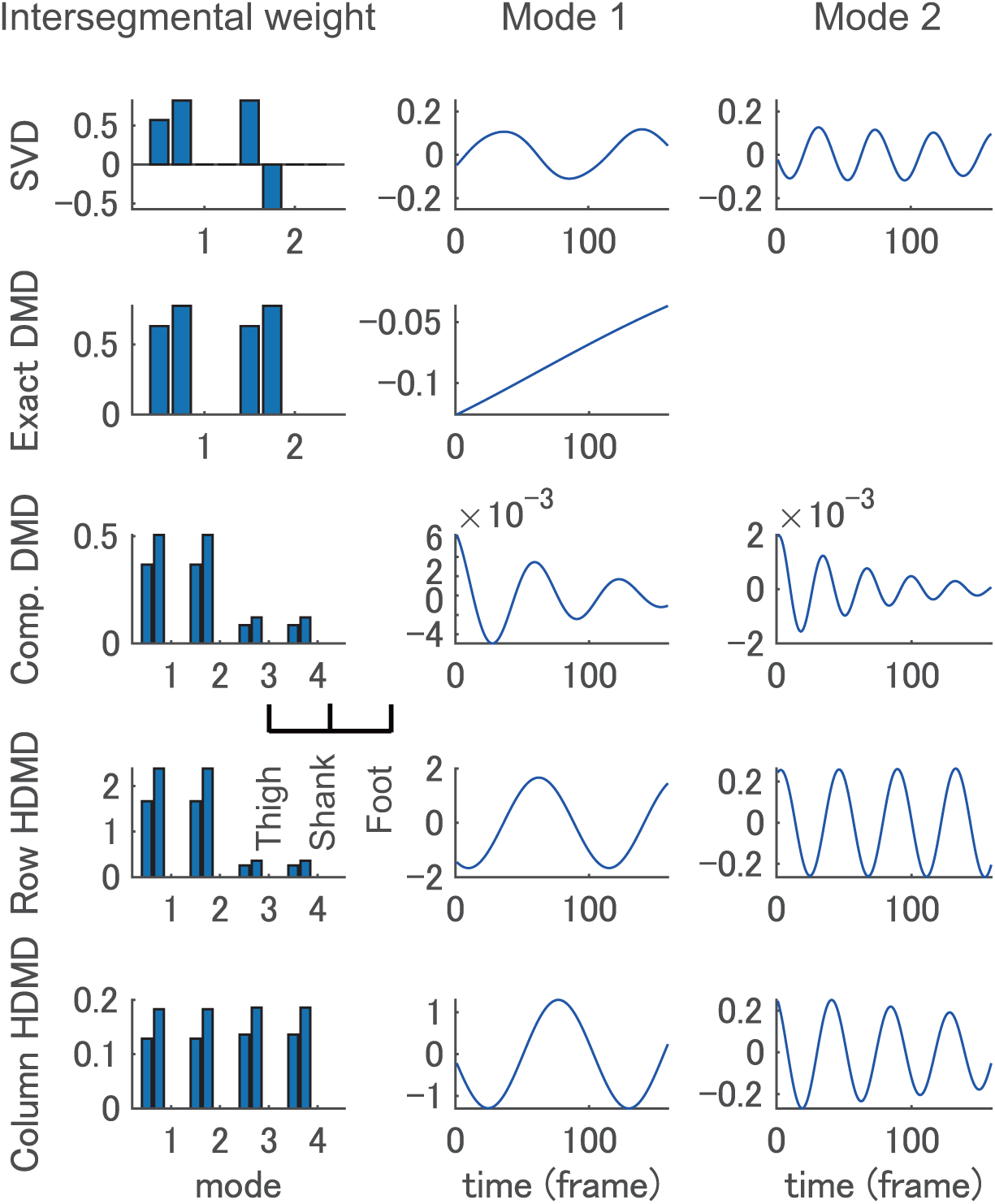
Decomposition of segmental angles of double pendulum simulation. Examples of decomposition results into intersegmental weights (or DMD modes) and time dynamics by various methods for double pendulum simulation are shown. Configurations are the same as in Fig 2, but we indicate two dominant modes because of the property of the system. Exact DMD and companion-matrix DMD show incorrect decomposition because of the smaller data dimension *d* = 2 (this is a well-known problem explained such as in [67]). Reconstruction error of the dominant modes is lower for SVD-based method (< 10^−15^), row-type Hankel DMD (0.0160 ± 0.0020), row-type Hankel DMD (0.0274 ± 0.0027), companion-matrix DMD (0.2566 ± 0.0050), and exact DMD (0.2663 ± 0.0216) in this order.

**Figure S4:**
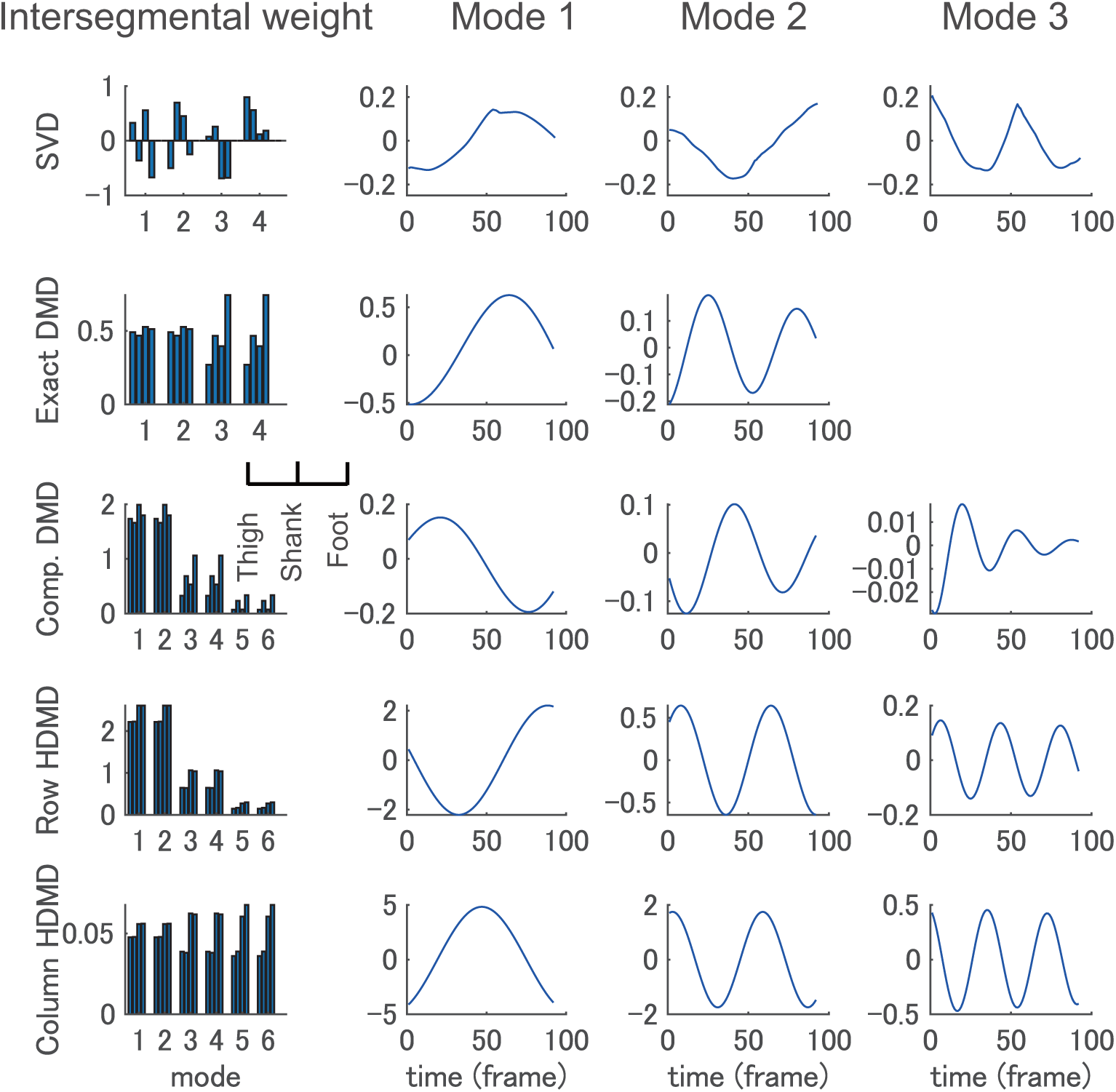
Decomposition of segmental angles of walking model simulation Examples of decomposition results into intersegmental weights (or DMD modes) and time dynamics by various methods for double pendulum simulation are shown. Configurations are the same as in Fig 2. Results were similar to Fig 2. Reconstruction error of the dominant modes are lower for SVD-based method (< 10^−15^), column-type Hankel DMD (0.0354 ± 0.0001), row-type Hankel DMD (0.0373 ± 0.0008), companion-matrix DMD (0.1356 ± 0.0348), and exact DMD (0.1411 ± 0.0043) in this order.

**Figure S5:**
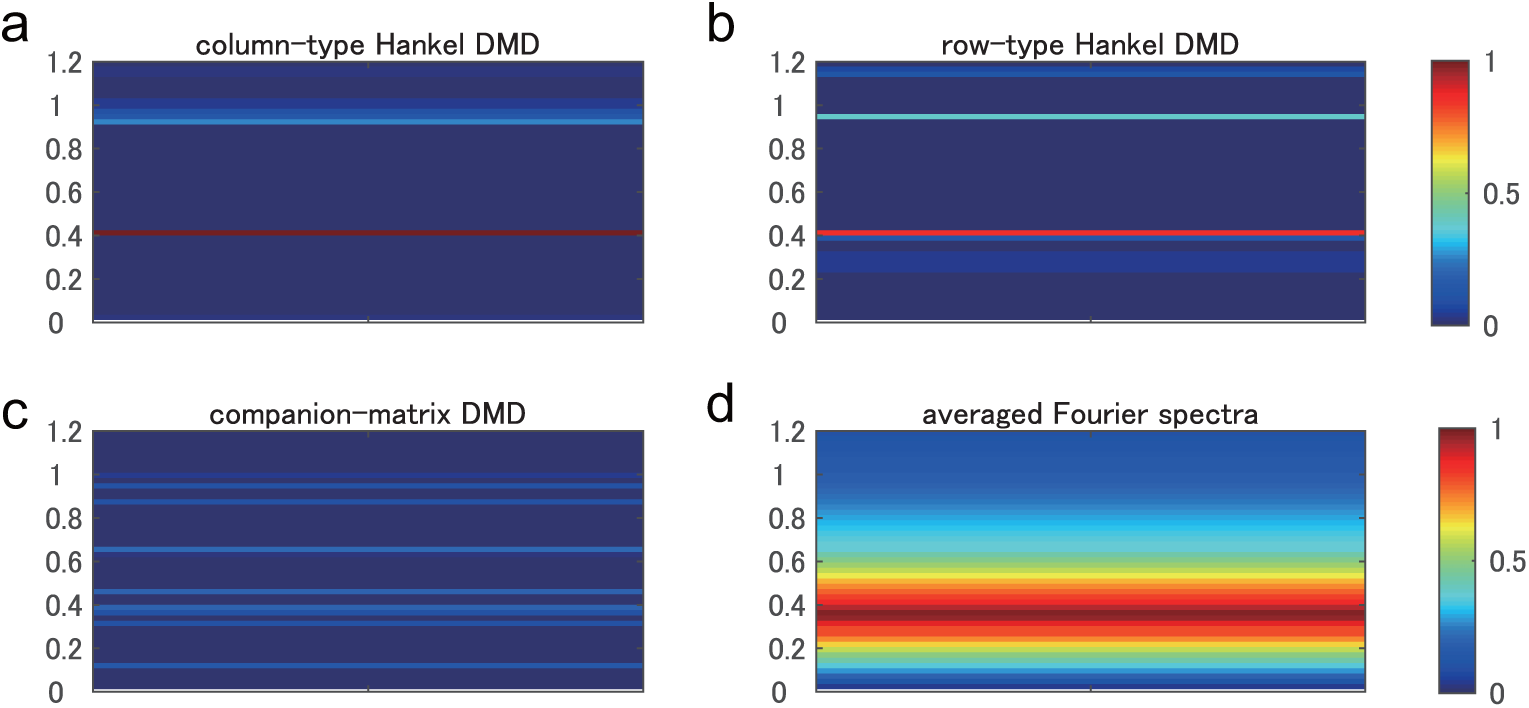
DMD eigenvalues of double pendulum simulation. Temporal frequency spectra in frequency domain to the gait frequency for (a) column-type and (b) row-type Hankel DMD, (c) companion-matrix DMD, and (d) Fourier transformation averaged among all participants for the double pendulum simulation data are shown. Configurations are the same as Fig 6 for clarity, but the horizontal axis indicates only one condition. Spectrum of each sequence was normalized so that the maximal norm is 1. Column- and row-type Hankel DMD seemed to obtain two eigenfrequencies (0.3815 and 0.9211 Hz). However, companion-matrix DMD did not distinctly indicate the eigenfrequencies. Averaged Fourier spectra were smoothed around only one eigenfrequency.

**Figure S6:**
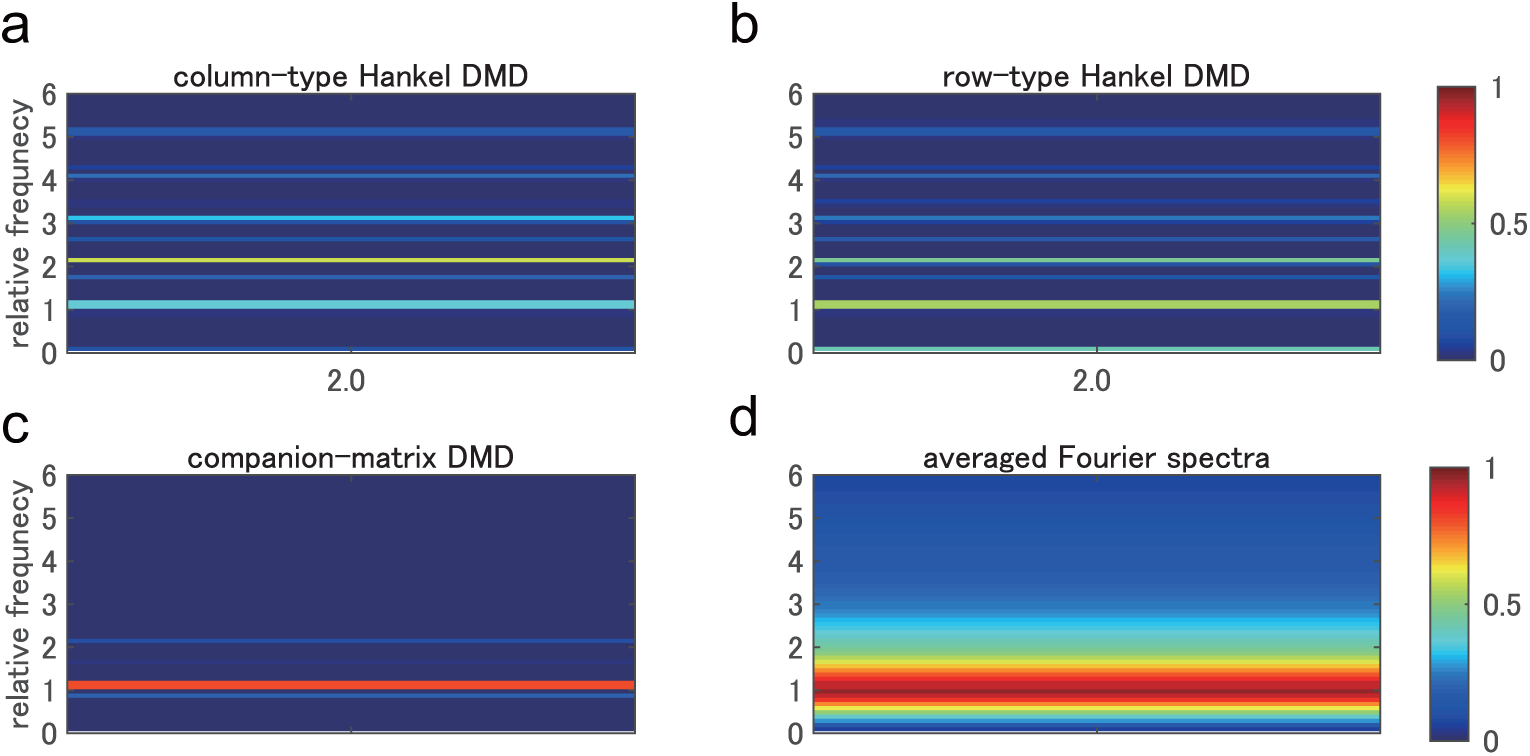
DMD eigenvalues of walking model simulation. Temporal frequency spectra in frequency domain to the gait frequency for (a) column-type and (b) row-type Hankel DMDs, (c) companion-matrix DMD, and (d) Fourier transformation averaged among all participants for the walking model simulation data are shown. Configurations are the same as in Fig 6 for clarity, but the horizontal axis indicates only one condition. Results were similar to those in Fig 6.

**Figure S7:**
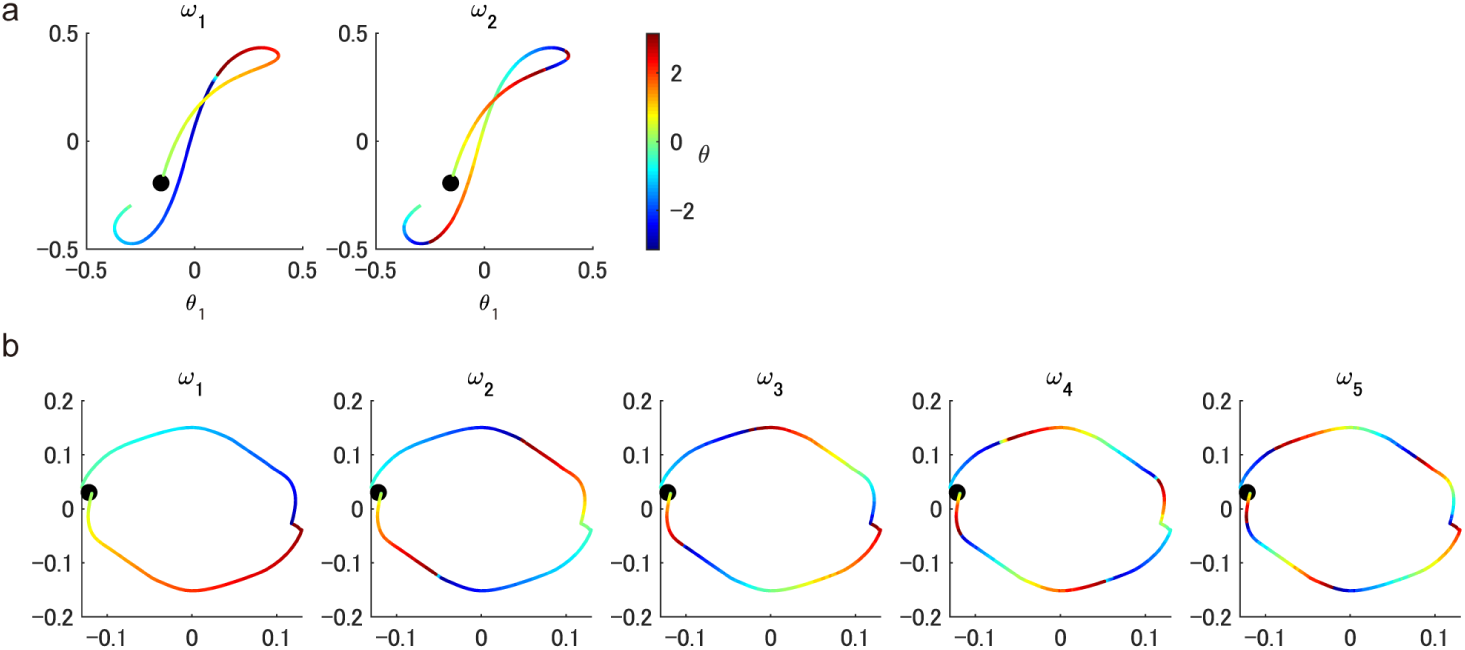
Phases of Koopman eigenfunctions for two simulation data. Phases computed by the argument of the Koopman eigenfunctions estimated by row-type Hankel DMD are shown for (a) the double pendulum simulation and (b) each walking speed and five harmonic frequencies. (a) For the pendulum simulation, because of the two-dimensional data, we visualize the phase on the two-dimensional trajectories. (b) For the walking simulation data, similarly to the human walking data, we visualize the phase on the two-dimensional structure obtained by the conventional SVD-based method. For clarity, we aligned the initial phases to near-zero values by the time-shift for the all eigenfunctions. All trajectories start from the black dots.

**Figure S8:**
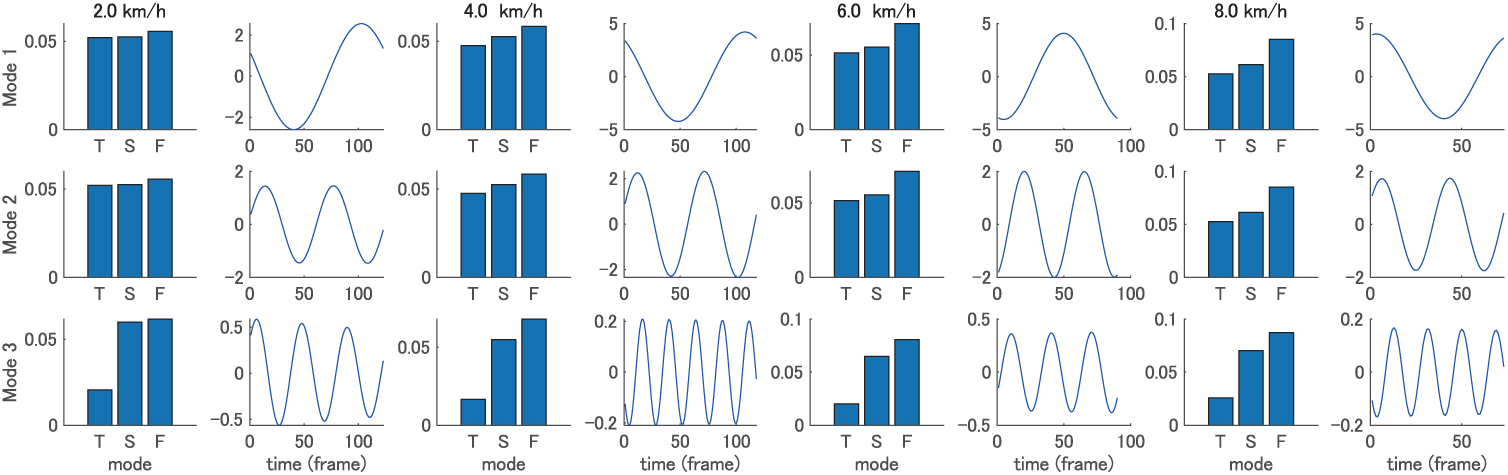
Representative examples of column-type Hankel DMD modes. Examples of three dominant column-type Hankel DMD modes and time dynamics during a 2.0 to 8.0 km/h walk are shown. Configurations are the same as in Fig 3. The DMD modes were similar among four walking speeds in terms of the larger weight of foot angle than that of thigh and shank angles.

## References

[1] N. Bernstein, The coordination and regulation of movement, Pergamon Press, London, 1967.

[2] R. Pfeifer, M. Lungarella, F. Iida, The challenges ahead for bio-inspired ‘soft’ robotics, Communications of the ACM 55 (2012) 76–87.

[3] C. Laschi, M. Cianchetti, Soft robotics: new perspectives for robot bodyware and control, Frontiers in Bioengineering and Biotechnology 2 (2014) 3.

[4] G. Taga, Y. Yamaguchi, H. Shimizu, Self-organized control of bipedal locomotion by neural oscillators in unpredictable environment, Biological Cybernetics 65 (3) (1991) 147–159.

[5] K. Fujii, Y. Yoshihara, H. Tanabe, Y. Yamamoto, Switching adaptability in human-inspired sidesteps: A minimal model, Frontiers in Human Neuroscience 11 (2017) 298.

[6] C. Fu, Y. Suzuki, K. Kiyono, P. Morasso, T. Nomura, An intermittent control model of flexible human gait using a stable manifold of saddle-type unstable limit cycle dynamics, Journal of the Royal Society Interface 11 (101) (2014) 20140958.

[7] T. P. Lillicrap, J. J. Hunt, A. Pritzel, N. Heess, T. Erez, Y. Tassa, D. Silver, D. Wierstra, Continuous control with deep reinforcement learning, arXiv preprint arXiv:1509.02971.

[8] S. Song, H. Geyer, A neural circuitry that emphasizes spinal feedback generates diverse behaviours of human locomotion, The Journal of Physiology 593 (16) (2015) 3493–3511.

[9] B. L. Davis, C. L. Vaughan, Phasic behavior of emg signals during gait: use of multivariate statistics, Journal of Electromyography and Kinesiology 3(1) (1993) 51–60.

[10] N. A. Borghese, L. Bianchi, F. Lacquaniti, Kinematic determinants of human locomotion., The Journal of Physiology 494(3) (1996) 863–879.

[11] B. J. West, N. Scafetta, Nonlinear dynamical model of human gait, Physical Review E 67 (5) (2003) 051917.

[12] V. B. Kokshenev, Dynamics of human walking at steady speeds, Physical Review Letters 93 (20) (2004) 208101.

[13] S. Shirasaka, W. Kurebayashi, H. Nakao, Phase reduction theory for hybrid nonlinear oscillators, Physical Review E 95 (1) (2017) 012212.

[14] M. E. Tinetti, S. F. Ginter, Identifying mobility dysfunctions in elderly patients: standard neuromuscular examination or direct assessment?, JAMA 259 (8) (1988) 1190–1193.

[15] D. C. Kerrigan, M. K. Todd, U. Della Croce, L. A. Lipsitz, J. J. Collins, Biomechanical gait alterations independent of speed in the healthy elderly: evidence for specific limiting impairments, Archives of Physical Medicine and Rehabilitation 79 (3) (1998) 317–322.

[16] N. Wenger, E. M. Moraud, J. Gandar, P. Musienko, M. Capogrosso, L. Baud, C. G. Le Goff, Q. Barraud, N. Pavlova, N. Dominici, et al., Spatiotemporal neuromodulation therapies engaging muscle synergies improve motor control after spinal cord injury, Nature Medicine 22 (2) (2016) 138.

[17] K. Fujii, S. Yoshioka, T. Isaka, M. Kouzaki, Unweighted state as a sidestep preparation improve the initiation and reaching performance for basketball players, Journal of Electromyography and Kinesiology 23 (6) (2013) 1467–1473.

[18] K. Fujii, M. Shinya, D. Yamashita, M. Kouzaki, S. Oda, Anticipation by basketball defenders: An explanation based on the three-dimensional inverted pendulum model, European Journal of Sport Science 14 (6) (2014) 538–546.

[19] F. Han, B. Reily, W. Hoff, H. Zhang, Space-time representation of people based on 3d skeletal data: A review, Computer Vision and Image Understanding 158 (2017) 85–105.

[20] M. Deng, C. Wang, F. Cheng, W. Zeng, Fusion of spatial-temporal and kinematic features for gait recognition with deterministic learning, Pattern Recognition 67 (2017) 186–200.

[21] F. Lacquaniti, R. Grasso, M. Zago, Motor patterns in walking, Physiology 14 (4) (1999) 168–174.

[22] G. Courtine, M. Schieppati, Tuning of a basic coordination pattern constructs straight-ahead and curved walking in humans, Journal of Neurophysiology 91 (4) (2004) 1524–1535.

[23] L. Bianchi, D. Angelini, G. Orani, F. Lacquaniti, Kinematic coordination in human gait: relation to mechanical energy cost, Journal of Neurophysiology 79 (4) (1998) 2155–2170.

[24] R. Grasso, L. Bianchi, F. Lacquaniti, Motor patterns for human gait: backward versus forward locomotion, Journal of Neurophysiology 80 (4) (1998) 1868–1885.

[25] Y. P. Ivanenko, R. Grasso, V. Macellari, F. Lacquaniti, Control of foot trajectory in human locomotion: role of ground contact forces in simulated reduced gravity, Journal of Neurophysiology 87 (6) (2002) 3070–3089.

[26] R. Poppele, G. Bosco, Sophisticated spinal contributions to motor control, Trends in Neurosciences 26 (5) (2003) 269–276.

[27] Y. P. Ivanenko, A. d’Avella, R. E. Poppele, F. Lacquaniti, On the origin of planar covariation of elevation angles during human locomotion, Journal of Neurophysiology 99 (4) (2008) 1890–1898.

[28] T. Funato, S. Aoi, N. Tomita, K. Tsuchiya, A system model that focuses on kinematic synergy for understanding human control structure, in: IEEE International Conference on Robotics and Biomimetics (ROBIO’12), IEEE, 2012, pp. 378–383.

[29] B. O. Koopman, Hamiltonian systems and transformation in hilbert space, Proceedings of the National Academy of Sciences 17 (5) (1931) 315–318.

[30] I. Mezić, Spectral properties of dynamical systems, model reduction and decompositions, Nonlinear Dynamics 41 (1) (2005) 309–325.

[31] M. W. Hirsch, S. Smale, R. L. Devaney, Differential equations, dynamical systems, and an introduction to chaos, Academic press, 2012.

[32] C. W. Rowley, I. Mezić, S. Bagheri, P. Schlatter, D. S. Henningson, Spectral analysis of nonlinear flows, Journal of Fluid Mechanics 641 (2009) 115–127.

[33] P. J. Schmid, Dynamic mode decomposition of numerical and experimental data, Journal of Fluid Mechanics 656 (2010) 5–28.

[34] J. L. Proctor, P. A. Eckhoff, Discovering dynamic patterns from infectious disease data using dynamic mode decomposition, International Health 7 (2) (2015) 139–145.

[35] B. W. Brunton, L. A. Johnson, J. G. Ojemann, J. N. Kutz, Extracting spatial-temporal coherent patterns in large-scale neural recordings using dynamic mode decomposition, Journal of Neuroscience Methods 258 (2016) 1–15.

[36] H. Arbabi, I. Mezić, Ergodic theory, dynamic mode decomposition, and computation of spectral properties of the koopman operator, SIAM Journal on Applied Dynamical Systems 16 (4) (2017) 2096–2126.

[37] H. Arbabi, I. Mezić, Study of dynamics in post-transient flows using koopman mode decomposition, Physical Review Fluids 2 (12) (2017) 124402.

[38] A. Mauroy, I. Mezić, On the use of fourier averages to compute the global isochrons of (quasi) periodic dynamics, Chaos: An Interdisciplinary Journal of Nonlinear Science 22 (3) (2012) 033112.

[39] I. Malkin, The methods of Lyapunov and Poincare in the theory of nonlinear oscillations, Gostexizdat, Moscow, 1949.

[40] A. T. Winfree, The geometry of biological time, Vol. 12, Springer Science & Business Media, 2001.

[41] D. Hansel, G. Mato, C. Meunier, Phase reduction and neural modeling, Concepts in Neuroscience 4 (1993) 192–210.

[42] E. Brown, J. Moehlis, P. Holmes, On the phase reduction and response dynamics of neural oscillator populations, Neural Computation 16 (4) (2004) 673–715.

[43] F. C. Hoppensteadt, E. M. Izhikevich, Weakly connected neural networks, Vol. 126, Springer Science & Business Media, 2012.

[44] S. R. Taylor, R. Gunawan, L. R. Petzold, F. J. Doyle III, Sensitivity measures for oscillating systems: Application to mammalian circadian gene network, IEEE Transactions on Automatic Control 53 (Special Issue) (2008) 177–188.

[45] P. Holmes, R. J. Full, D. Koditschek, J. Guckenheimer, The dynamics of legged locomotion: Models, analyses, and challenges, SIAM Review 48 (2) (2006) 207–304.

[46] K. Fujii, T. Kawasaki, Y. Inaba, Y. Kawahara, Prediction and classification in equation-free collective motion dynamics, PLoS Computational Biology 14 (11) (2018) e1006545.

[47] T. Nomura, K. Kawa, Y. Suzuki, M. Nakanishi, T. Yamasaki, Dynamic stability and phase resetting during biped gait, Chaos: An Interdisciplinary Journal of Nonlinear Science 19 (2) (2009) 026103.

[48] J. A. Nessler, T. Spargo, A. Craig-Jones, J. G. Milton, Phase resetting behavior in human gait is influenced by treadmill walking speed, Gait & Posture 43 (2016) 187–191.

[49] J. A. Nessler, S. Heredia, J. Bélair, J. Milton, Walking on a vertically oscillating treadmill: phase synchronization and gait kinematics, PLoS One 12 (1) (2017) e0169924.

[50] T. Funato, Y. Yamamoto, S. Aoi, T. Imai, T. Aoyagi, N. Tomita, K. Tsuchiya, Evaluation of the phase-dependent rhythm control of human walking using phase response curves, PLoS Computational Biology 12 (5) (2016) e1004950.

[51] Y. P. Ivanenko, G. Cappellini, N. Dominici, R. E. Poppele, F. Lacquaniti, Modular control of limb movements during human locomotion, Journal of Neuroscience 27 (41) (2007) 11149–11161.

[52] F. Hebenstreit, A. Leibold, S. Krinner, G. Welsch, M. Lochmann, B. M. Eskofier, Effect of walking speed on gait sub phase durations, Human Movement Science 43 (2015) 118–124.

[53] B. Kibushi, S. Hagio, T. Moritani, M. Kouzaki, Speed-dependent modulation of muscle activity based on muscle synergies during treadmill walking, Frontiers in Human Neuroscience 12 (2018) 4.

[54] B. Kibushi, T. Moritani, M. Kouzaki, Local dynamic stability in temporal pattern of intersegmental coordination during various stride time and stride length combinations, Experimental Brain Research (2018) 1–15.

[55] E. Attinger, A. Anne, D. McDonald, Use of fourier series for the analysis of biological systems, Biophysical Journal 6 (3) (1966) 291.

[56] J. M. Brownjohn, A. Pavic, P. Omenzetter, A spectral density approach for modelling continuous vertical forces on pedestrian structures due to walking, Canadian Journal of Civil Engineering 31 (1) (2004) 65–77.

[57] G. E. Grossman, R. J. Leigh, L. Abel, D. J. Lanska, S. Thurston, Frequency and velocity of rotational head perturbations during locomotion, Experimental Brain Research 70 (3) (1988) 470–476.

[58] D. A. Winter, H. G. Sidwall, D. A. Hobson, Measurement and reduction of noise in kinematics of locomotion, Journal of Biomechanics 7 (2) (1974) 157–159.

[59] T. Funato, S. Aoi, H. Oshima, K. Tsuchiya, Variant and invariant patterns embedded in human locomotion through whole body kinematic coordination, Experimental Brain Research 205 (4) (2010) 497–511.

[60] S. L. Brunton, J. L. Proctor, J. N. Kutz, Discovering governing equations from data by sparse identification of nonlinear dynamical systems, Proceedings of the National Academy of Sciences 201517384.

[61] A. Barliya, L. Omlor, M. A. Giese, T. Flash, An analytical formulation of the law of intersegmental coordination during human locomotion, Experimental Brain Research 193 (3) (2009) 371–385.

[62] K. Fujii, T. Isaka, M. Kouzaki, Y. Yamamoto, Mutual and asynchronous anticipation and action in sports as globally competitive and locally coordinative dynamics, Scientific Reports 5. doi:10.1038/srep16140.

[63] K. Fujii, S. Yoshioka, T. Isaka, M. Kouzaki, The preparatory state of ground reaction forces in defending against a dribbler in a basketball 1-on-1 dribble subphase, Sports Biomechanics 14 (1) (2015) 28–44.

[64] L. Hak, H. Houdijk, P. J. Beek, J. H. van Dieën, Steps to take to enhance gait stability: the effect of stride frequency, stride length, and walking speed on local dynamic stability and margins of stability, PLoS One 8 (12) (2013) e82842.

[65] G. Torres-Oviedo, J. M. Macpherson, L. H. Ting, Muscle synergy organization is robust across a variety of postural perturbations, Journal of Neurophysiology 96 (3) (2006) 1530–1546.

[66] J. H. Tu, C. W. Rowley, D. M. Luchtenburg, S. L. Brunton, J. N. Kutz, On dynamic mode decomposition: Theory and applications., Journal of Computational Dynamics 1 (2) (2014) 391–421.

[67] J. N. Kutz, S. L. Brunton, B. W. Brunton, J. L. Proctor, Dynamic Mode Decomposition: Data-Driven Modeling of Complex Systems, SIAM, 2016.

[68] M. O. Williams, I. G. Kevrekidis, C. W. Rowley, A data-driven approximation of the koopman operator: Extending dynamic mode decomposition, Journal of Nonlinear Science 25 (6) (2015) 1307–1346.

[69] Y. Kawahara, Dynamic mode decomposition with reproducing kernels for koopman spectral analysis, in: Advances in Neural Information Processing Systems 29, 2016, pp. 911–919.

[70] N. Takeishi, Y. Kawahara, T. Yairi, Learning koopman invariant subspaces for dynamic mode decomposition, in: Advances in Neural Information Processing Systems 30, 2017, pp. 1130–1140.

[71] K. K. Chen, J. H. Tu, C. W. Rowley, Variants of dynamic mode decomposition: boundary condition, koopman, and fourier analyses, Journal of Nonlinear Science 22 (6) (2012) 887–915.

[72] S. L. Brunton, B. W. Brunton, J. L. Proctor, E. Kaiser, J. N. Kutz, Chaos as an intermittently forced linear system, Nature Communications 8 (1) (2017) 19.

[73] M. Gavish, D. L. Donoho, The optimal hard threshold for singular values is 4*/√*3, IEEE Transactions on Information Theory 60 (8) (2014) 5040–5053.

[74] E. Bradley, H. Kantz, Nonlinear time-series analysis revisited, Chaos: An Interdisciplinary Journal of Nonlinear Science 25 (9) (2015) 097610.

[75] M. B. Kennel, R. Brown, H. D. Abarbanel, Determining embedding dimension for phase-space reconstruction using a geometrical construction, Physical Review A 45 (6) (1992) 3403.

[76] J. A. Nessler, C. J. De Leone, S. Gilliland, Nonlinear time series analysis of knee and ankle kinematics during side by side treadmill walking, Chaos: An Interdisciplinary Journal of Nonlinear Science 19 (2) (2009) 026104.

[77] N. Mangan, J. Kutz, S. Brunton, J. Proctor, Model selection for dynamical systems via sparse regression and information criteria, Proceedings of the Royal Society A: Mathematical, Physical and Engineering Sciences 473 (2204).

[78] H. Akaike, A new look at the statistical model identification, IEEE Transactions on Automatic Control 19 (6) (1974) 716–723.

